# Reinforcement learning enhances training efficiency in high-performance athletes

**DOI:** 10.1101/2024.04.22.590558

**Authors:** Christian Magelssen, Matthias Gilgien, Simen Leithe Tajet, Thomas Losnegard, Per Haugen, Robert Reid, Romy Frömer

## Abstract

Skilled athletes need powerful movement strategies to solve tasks effectively. Typically, athletes learn these strategies with instruction-based teaching methods where coaches offer athletes a correct solution. Inspired by recent evidence from decision neuroscience, we asked whether skilled athletes learn strategy choices better with an evaluation-based training strategy (reinforcement learning). To address this question, we conducted a three-day learning experiment with skilled alpine ski racers (n=98) designed to improve their performance on flat slopes on slalom with four strategies at their disposal to achieve this goal. We compared performance and strategy choices of three groups: a reinforcement learning group, that only received feedback about their race times after every run, a supervised (free choice) learning group, that received strategy instructions from their coach, and a supervised (target skill) learning group, being coached to use the theoretically optimal strategy for skiing well on flats. We found that despite making similar strategy choices, the skiers in the reinforcement learning group, showed greater improvements in their race times during the training sessions than their counterparts in the supervised (free choice) learning group and outperformed them during a subsequent retention test. Surprisingly, the skiers in the reinforcement learning group even showed descriptively (but not significantly) better performance than those in the supervised (target skill) learning group. Our findings show that reinforcement learning can be an effective training strategy for improving strategy choices and performance among skilled athletes, even among the best ones.

## 1 Introduction

Skilled performance involves precise and accurate execution of well-chosen strategies[1–10]. For many athletes (even the most skilled ones), these strategies may not always be optimal and can hold them back from unleashing their full potential. In those cases, athletes may need to try new strategies and learn their values to achieve further improvements. Unfortunately, many athletes are stuck on plateaus because they lack knowledge of better alternatives and rely on conventional teaching methods that focus on refining existing strategies [11–15]. Are there better training methods that can improve elite performance, by teaching athletes new strategies’ values in more efficient ways than conventional teaching? If so, these training methods could greatly impact the training of present an d future learners.

In contemporary teaching practice, athletes commonly learn which movement strategy to execute with instruction-based teaching. That is, a coach or teacher advises the athlete ‘what to do’ (e.g., suggesting taking a shorter line around the gate) followed by corrective feedback (e.g., encouraging the learner to further shorten the line) [16–18]. This teaching strategy can be likened to supervised learning in motor learning, where the difference between the desired strategy and the athlete’s movement outcome represents a teaching signal for skill improvement [2, 19, 20]. Through practice, this teaching signal can bring the athlete closer to executing what is assumed to be the correct choice. But is this teaching strategy truly the most effective approach for enabling learners to select and employ effective strategies?

Although supervised learning serves as a reasonable approach to improve athletes’ strategies, the teaching strategy has limitations that may prevent athletes from converging on an optimal strategy. First, the coach’s advice may not be the optimal solution, as what coaches judge as a good strategy does not always align with reality, even for the best-trained eye [21, 22]. Athletes might therefore miss opportunities to discover the best strategy when coaches opt for suboptimal strategies [12, 13]. Worse, these learned suboptimal strategies might turn into habits that can be difficult to break [23]. Second, athletes trained through supervised learning might also be constrained to adopting a single (’universal’) strategy for all situations rather than acquiring a repertoire of strategies and discerning the most effective strategies for each specific scenario. Finally, it remains uncertain whether the prescriptive approach is the most effective teaching strategy for enabling athletes to achieve long-lasting learning effects [16, 17, 24, 25].

Athletes can also learn to choose effective strategies without depending on direct advice from a coach. The cornerstone of reinforcement learning [26] is that athletes can learn by exploring strategies and evaluating their outcomes, using the successes and failures of outcomes as teaching signals. That is, the athlete learns the value of different strategies which allows them to finally pick the best solution, rather than being told the putatively correct solution to the problem. Specifically, these values are learned by comparing a given choice’s outcomes with the choice’s currently expected outcome. Outcomes that exceed or fall short of expectations result in errors in reward prediction, signaling that the learner must update their predictions to better anticipate future rewards following that action [27]. These reward prediction errors are then incorporated to form a new and better estimate of expected reward by updating expectations through a weighted running average. The computations underlying this form of value-learning of choices have been tremendously powerful in explaining human and animal learning [28–33] as well as training AI to perform complex tasks such as computer games starting from pixel inputs, only[34]. In addition, reinforcement learning has been shown to improve motor skill learning on simple tasks performed in the laboratory [35–38]. Based on this evidence, we asked whether reinforcement learning offers a better alternative for training learners to make better decisions about strategies than traditional supervised learning with a coach.

To address this question, we conducted a three-day learning experiment with ninety-eight skilled alpine ski racers. To achieve performance improvement among this skilled cohort of athletes, we focused on improving flat sections in slalom, an area with considerable potential for enhancement, even among the best skiers[39], and delineated four strategies, each carefully selected to enhance performance in this section by taking advantage of principles from mechanics (Fig. 2a). To study how different instructions and feedback influenced the learning and choices of these strategies, we assigned the skiers to three different learning groups (Fig. 2b). In the supervised (target skill) learning group, we recruited ski coaches to instruct skiers to select the strategy that we defined as the theoretically best strategy based on computational modeling [40–42] and field observations of elite skiers [43, 44]. This group serves as a benchmark for the upper limit of performance achievable through optimal strategy choices and thus offers an upward comparison for the performance yielded through reinforcement learning of strategy selection. In the supervised (free choice) learning group, we recruited ski coaches from the tested ski teams to train them on the strategy they believed to be the best or most appropriate for the skier. Coaches in the two supervised learning groups saw the skiers’ times but were instructed to not disclose these to the skiers. In contrast, in the reinforcement learning group, the skiers chose a strategy on every trial and saw their race times immediately after each trial to inform them about which strategy to select. We hypothesized that the skiers in the reinforcement learning group would learn to choose better strategies and thus achieve better performance than skiers subject to traditional supervised learning with a coach (supervised learning: free choice), possibly reaching similar performance as the supervised (target skill) learning group that uses the best strategy throughout.

## 2 Method

### 2.1 Participants

Conducting studies on alpine ski racing poses challenges related to environmental control and resource constraints. Our sample size approach was justified by the ‘resource constraints’ and the ‘whole population’ criteria [45] and involved recruiting as many skiers as possible during June 2023, when we had a short time window to test skiers in the indoor ski hall. We set the minimum sample size to 80 skiers, which we deemed appropriate for this context. Prior to data collection, data and power simulations for sample sizes of 80, 100, and 120 skiers were conducted, revealing simulated powers of 0.60, 0.75, and 0.80, respectively, for the smallest effect size of interest (0.3 second difference between groups) (https://osf.io/c4t28). The smallest effect size of interest was based on our knowledge of alpine skiing and discussions with coaches, but it was intended for a 50-meter longer course and more training sessions than we ultimately ended up using due to practical considerations. We deliberately opted to recruit skiers with diverse skill levels for the study to augment the generalizability of our findings. However, to ensure a sufficient skill level to handle the specific icy snow conditions prepared in the ski hall, we recruited only skiers aged 15 and older. The sample size justification, task design, and analysis plan were preregistered before data collection (https://osf.io/tfb2w).

We managed to recruit ten alpine ski teams comprising 98 alpine ski racers from Norway and Sweden (age M = 18.1 years, SD= 2; 40 females, 58 males). Two skiers were excluded from the analysis due to an injury prior to the study (n=1) or sickness during the study (n=1); thus, a total of 96 skiers completed the entire study and were included in the analysis. Among the ski groups tested were five ski academies, three senior development teams, and two national ski teams. These skiers were generally highly skilled, with a median world rank of 605, but there was also considerable variability, as indicated by a substantial interquartile range (Q1 = 248, Q3 = 1390.5). A smaller subset of the participants (n = 13) were not world-ranked, as they had yet to compete in internationally sanctioned races that form the basis for calculating athlete points and rankings. Table 1 provides demographic information for each learning group.

**Table 1.**
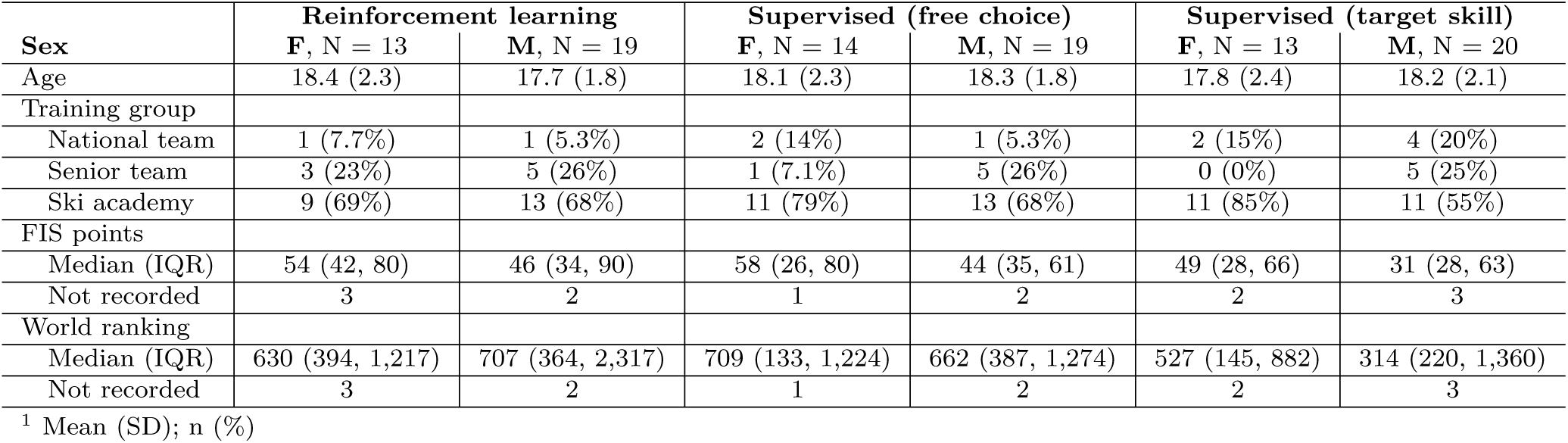
Skier characteristics for each learning group.

| Sex | Reinforcement learning |  | Supervised (free choice) |  | Supervised (target skill) |  |
| --- | --- | --- | --- | --- | --- | --- |
|  | F, N = 13 | M, N = 19 | F, N = 14 | M, N = 19 | F, N = 13 | M, N = 20 |
| Age | 18.4 (2.3) | 17.7 (1.8) | 18.1 (2.3) | 18.3 (1.8) | 17.8 (2.4) | 18.2 (2.1) |
| Training group |  |  |  |  |  |  |
| National team | 1 (7.7%) | 1 (5.3%) | 2 (14%) | 1 (5.3%) | 2 (15%) | 4 (20%) |
| Senior team | 3 (23%) | 5 (26%) | 1 (7.1%) | 5 (26%) | 0 (0%) | 5 (25%) |
| Ski academy | 9 (69%) | 13 (68%) | 11 (79%) | 13 (68%) | 11 (85%) | 11 (55%) |
| FIS points |  |  |  |  |  |  |
| Median (IQR) | 54 (42, 80) | 46 (34, 90) | 58 (26, 80) | 44 (35, 61) | 49 (28, 66) | 31 (28, 63) |
| Not recorded | 3 | 2 | 1 | 2 | 2 | 3 |
| World ranking |  |  |  |  |  |  |
| Median (IQR) | 630 (394, 1,217) | 707 (364, 2,317) | 709 (133, 1,224) | 662 (387, 1,274) | 527 (145, 882) | 314 (220, 1,360) |
| Not recorded | 3 | 2 | 1 | 2 | 2 | 3 |

<sup>1</sup> Mean (SD); n (%)

In addition, we recruited coaches for the two supervised learning groups. This sampling process was carried out pragmatically due to space and time constraints in the ski hall, which made it impossible to test all ski teams simultaneously. The 10 ski teams with alpine skiers that were recruited were therefore divided into 4 groups for which the study was conducted at different times. For each of the four groups, we recruited two coaches from the ski teams to serve as coaches in the supervised (free choice) learning group, totaling 8 coaches (2 women; 6 men). These coaches had extensive coaching experience in coaching alpine ski racers. In addition, for each group of ski teams that completed the experiment together, we recruited a third coach to instruct the skiers in the supervised (target skill) learning group. To ensure that these coaches had sufficient credibility to make the skiers buy into our theoretical best strategy, we selectively recruited three highly experienced coaches from the Norwegian alpine ski team (one coach served twice). Importantly, all the coaches remained unaware of the experimental manipulation. Table 2 provides demographic information about the coaches. All the skiers and coaches provided informed consent before the study. The study was approved by the Human Research Ethics Committee of The Norwegian School of Sport Sciences (ref. 279-040523) and the Norwegian Agency for Shared Services in Education and Research (ref. 871468), and was performed in accordance with their guidelines and regulations.

**Table 2.**
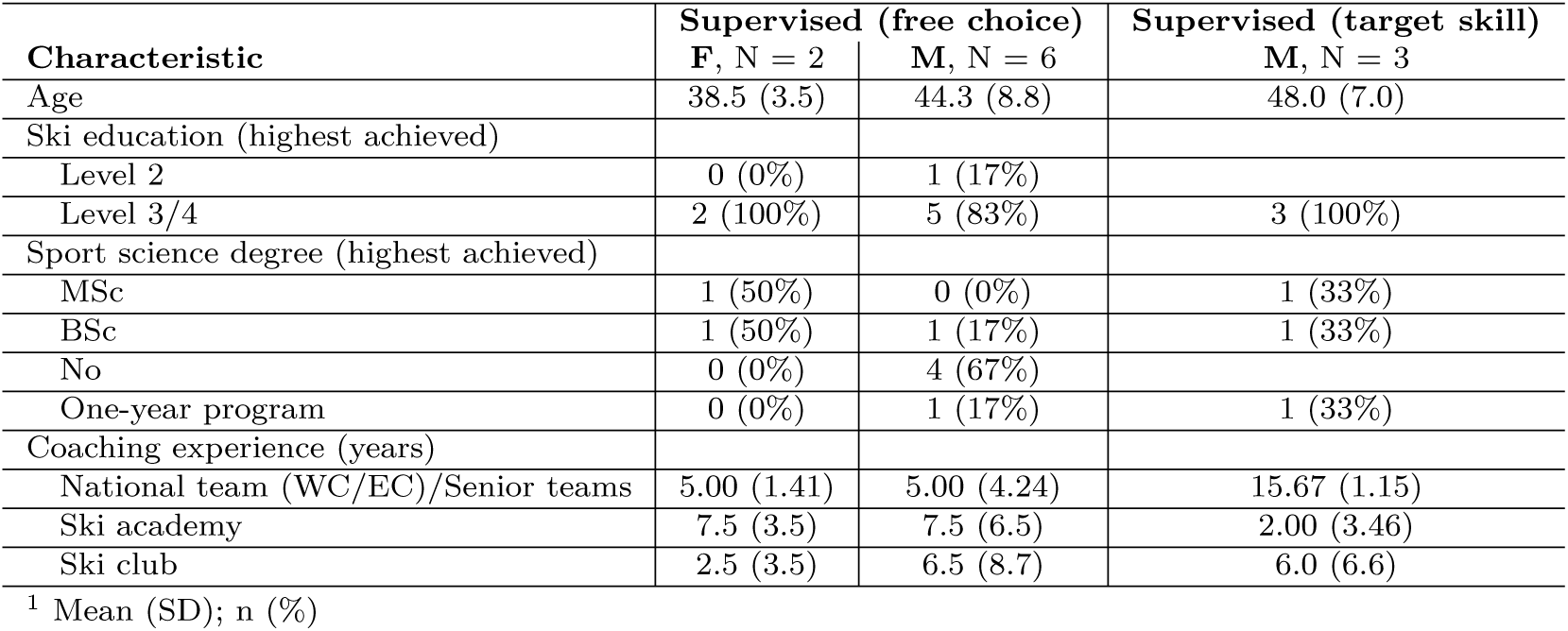
Coach characteristics for each learning group.

### 2.2 The setup

The experiment was conducted in the indoor ski hall SNØ in Oslo, Norway (https://snooslo.no/). In this hall, we used a 210-meter-long flat slope section of the race hill, which we water-injected before testing each group of skiers to ensure uniform and fair snow conditions for all skiers (Supplementary Methods A for detailed description). With our chosen course setup, this 210-meter-long flat section allowed space for 19 slalom gates.

We used two types of slalom courses in the experiment (Fig. 1a). The main slalom course was used in all sessions, except during the transfer test, and featured a 10 m distance and a 1.9 m offset. The course distance aligned with our previous study [44], but we opted for a slightly larger offset to better suit the skill level of our skiers. The transfer test evaluated how well the skiers transferred their learning to a new slalom course more realistic to a typical alpine ski race course. To assess this, we set a course with a progression in gate offset, starting with five gates at a 2.2-meter offset, followed by seven gates at a 1.7-meter offset, and concluding with seven gates at a 1.2-meter offset. Although we did not expect radical differences in strategy effects, we anticipated a greater emphasis on rocking the skis forward in gates with a 2.2-meter offset than in those with a 1.2-meter offset to enhance turn exit release. Both courses were set with stubbies (short gates) instead of long gates to minimize energy dissipation upon hitting the gate [46]. Using long gates can also be a distracting element in that skiers’ attention is allocated to clearing the gate instead of focusing on executing the skill. Finally, this approach helped us avoid creating holes in the course, which can occur when the long gate is forcefully slammed into the ground. To minimize wear and tear on the course, we set two parallel and identical courses and routinely shifted between them. See Supplementary Methods B for a detailed explanation of our course setting procedure.

**Fig. 1.**
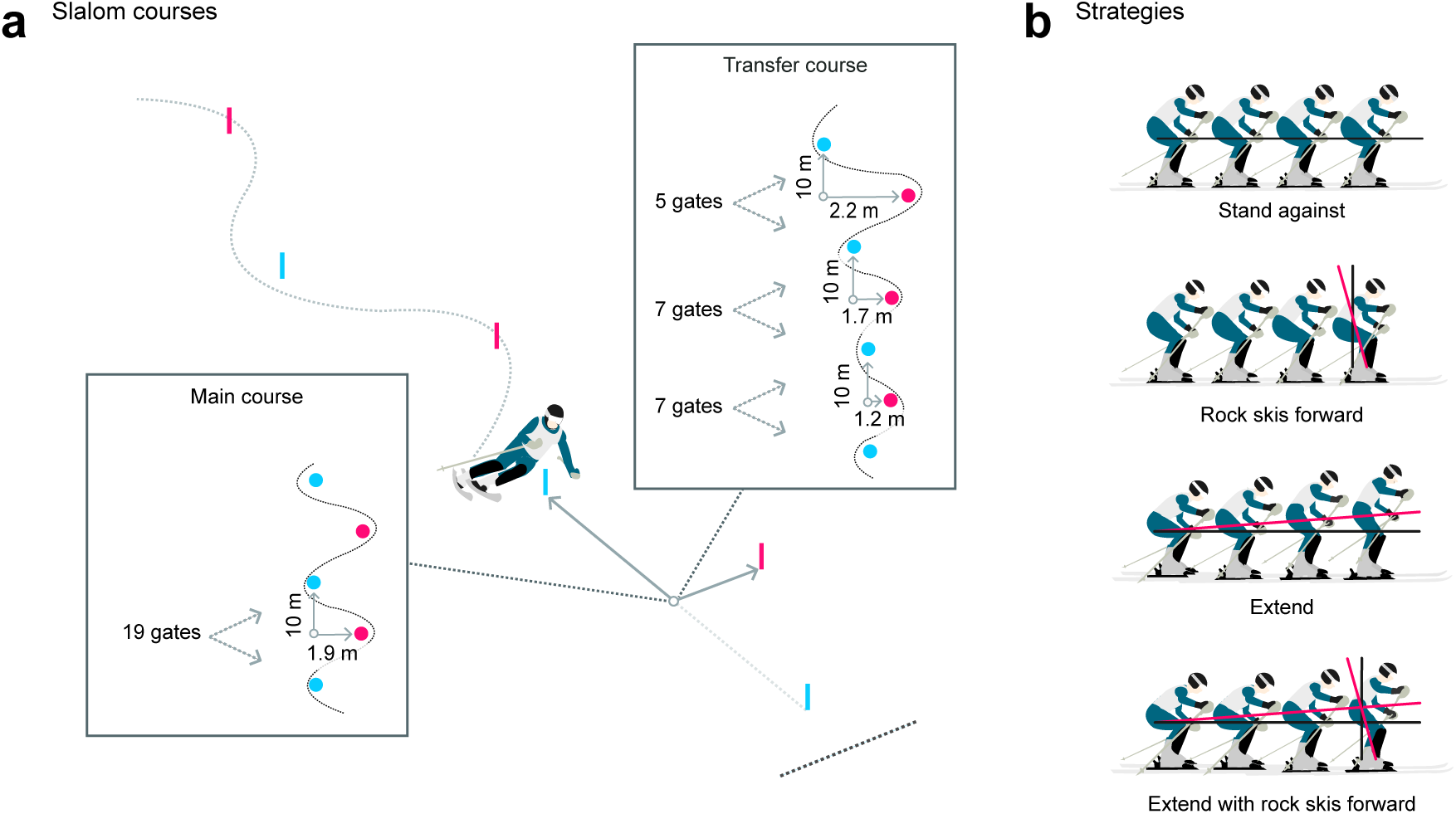
**a.** Illustrations of the two slalom courses used in the study. The main slalom course was a rhythmic course deployed in all sessions except for the transfer session. The course setting for the transfer session involved a progression in gate offset, starting with the largest offset and ending with the smallest offset. **b.** Illustration of the strategies defined to enhance racing performance on flat terrain in slalom: The ”stand against” strategy emphasized maintaining a stable stance against external forces without body extension along the body’s longitudinal axis or rocking skis forward; ”Rock skis forward” involved rocking skis forward from gate passage to completion of the turn; The ”extend” strategy involves extending the body from a laterally tilted position during the turn, closer to the turn’s center of rotation; The ”extend with rocking skis forward” was expected to be the best strategy combining the two effects from extending and rocking skis forward, and we therefore defined this as the theoretical best strategy

The start gate was positioned 20 meters before the first gate. Skiers were required to start in a static position to ensure consistency in the starts, with the toe piece of the binding placed behind the starting gate. The skiers started by putting their skis in parallel and lifting the poles without using poling or skating for propulsion (Supplementary Video illustrates the starting procedure and setup). We recorded the times using a wireless photocell timing system (HC Timing wiNode and wiTimer; Oslo, Norway). Timing started when the skier crossed the first photocell pair situated 10 meters below the starting gate.

### 2.3 Experimental design

We employed a between-subjects design and posed the learning question of discovering effective strategies as an *n*-armed bandit problem [26]. The essence of this problem is that a learner repeatedly tries different options and observes their outcomes to learn which strategy is the best and, therefore, which one to choose. Finding the best strategy requires a delicate balance between exploiting the strategy known to yield the best payoff and exploring alternative strategies that may offer superior benefits. In our study, the options consisted of four strategies that skiers could employ to improve their race times on flat slopes in slalom, grounded in physics-based coaching manuals for alpine ski racing [40, 41, 47, 48], biomechanical research on elite skiers [44, 49–51] and common strategies used by coaches. The four strategies were named ”stand against”, ”rock skis forward”, ”extend”, and ”extend with rock skis forward (see Fig. 1b for a strategy illustration and Supplementary Methods C for an extended explanation). To study how instruction and feedback drove learning to select effective strategies, we designed and allocated skiers to three learning groups, which allowed us to compare reinforcement learning with traditional supervised learning with a coach: For the supervised (target skill) learning group, we provided the best possible training program by engaging highly experienced coaches who explained to the skiers that the ‘extend with rock skis forward’ strategy was the most effective for skiing fast on flat terrain in slalom, citing evidence from research literature in alpine skiing mechanics [40, 41, 49]. The coach then instructed the skiers to adopt this strategy and provided feedback on its execution after each trial. Note that the coach had access to the skiers after each trial but was prohibited from communicating the race time with the skiers.

In the supervised (free choice) learning group, the skiers were assigned to two coaches recruited from the tested group of ski teams. To balance the skiers’ skill levels between the two coaches, we created new blocks from the ranked list from baseline testing and randomly assigned them to the coaches. We instructed the coaches to improve the skiers’ race times as much as possible with our four defined strategies to their disposal to achieve this goal. For each trial, the coach selected a strategy for the skier, observed the skier during the trial and provided feedback on its execution afterward. Similar to the supervised (target skill) learning group, the coaches had access to the skiers’ data after each trial but could not share this information with the skiers.

In contrast, the reinforcement learning group was not assigned to any coach. Instead of having a coach deciding the skiing strategy for them, the skiers in this learning group were told to choose a strategy for each trial by themselves to ski the course as fast as possible. To help the skiers choose and learn from evaluation, this group could see their racing times immediately after they crossed the finish line. Although this group had no coach, we assigned a person to communicate with the skiers to record their choices and encourage them to try skiing quickly to prevent boredom effects.

### 2.4 Procedures

Fig. 2a illustrates the procedures employed in the study. In the baseline test, the skiers began with two warmup runs: one in a free skiing warmup course and one in a specific warm-up in the slalom course. During these warm-up runs, skiers were instructed, trained and verified on the start procedure by an instructor. As a first run in the baseline assessment, skiers completed a straight-gliding run, where they skied straight down from start to finish in a static, upright slalom posture. Subsequently, the skiers completed four runs in the slalom course. Skiers were encouraged to ski as quickly as they could, but they could not see their times, nor did they receive any instructions on how to perform well.

**Fig. 2.**
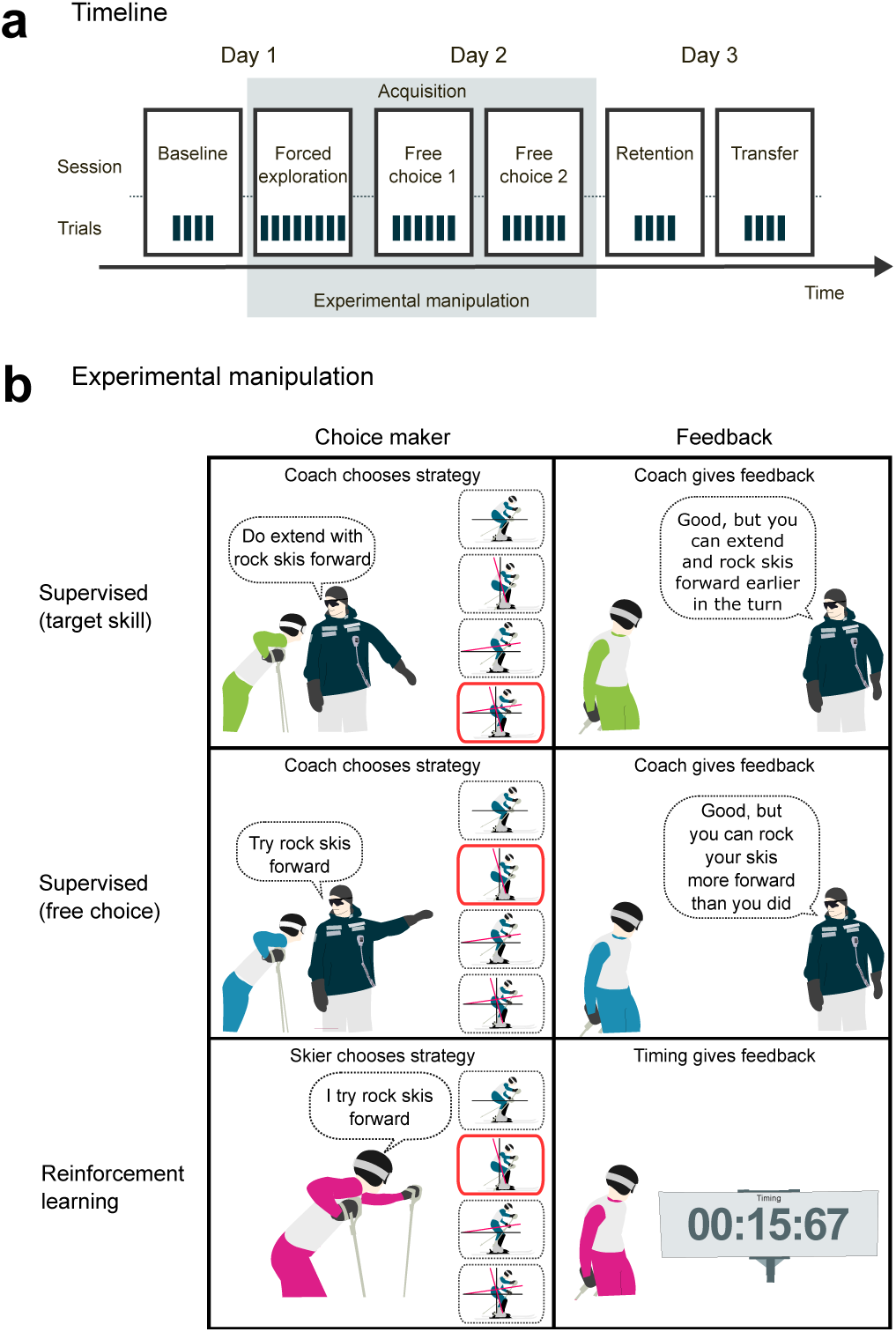
Illustration of the experimental design and procedure. **a.** Timeline of the three-day learning experiment. During the baseline session, the skiers raced a slalom course in the shortest time possible without receiving race time feedback. The skiers were then assigned to one of three learning groups (see b). In their assigned group, the skiers underwent three acquisition sessions, comprising one forced exploration session (skiers performed all strategies) and two free choice sessions (skiers or coaches could choose strategies themselves). On the last day, skiers completed a retention and transfer test where they could pick strategies themselves, again without receiving race time feedback. **b.** Illustration of the learning groups in the study. The supervised (target skill) learning group involved coaches consistently choosing the theoretically best strategy (except during forced exploration), while in the supervised (free choice) learning group coaches freely selected strategies. The skiers in both of these learning groups received feedback on strategy execution from their assigned coach, while the skiers in the reinforcement learning group independently selected strategies and received feedback from the timing system to facilitate value learning of each strategy.

After the initial baseline assessment, the skiers took a 60-minute break. In the meantime, we allocated the skiers to the three learning groups deploying a randomized-blocked approach to account for pre-existing differences in the skiers’ performance levels [52]. Specifically, we computed each skier’s average across the four trials at baseline and ranked them accordingly. We then created *n* blocks with block sizes corresponding to our three learning groups for the entire list of skiers and assigned these skiers to these predefined blocks. Finally, we randomly allocated the skiers to the different learning groups within each block (Fig. 2b).

The learning groups participated in sessions at different times to prevent treatment diffusion [52]. As the ski group comprised teams of skiers who knew each other well and who resided together, we explicitly emphasized the importance of keeping information about the sessions private. To stay within the time frame at the ski hall, the two supervised learning groups underwent training together. To facilitate this, we arranged stations in the finishing area with space and vision dividers and ample space to impede communication between coaches in the supervised learning groups (Supplementary Methods D). In addition, we developed a Python script that fetched the race times from the timing system, filtered the times for each coach, and transmitted them to the station where the coach was located, ensuring that no information leaked between the coaches. The learning group (reinforcement learning versus supervised learning) that initiated after the baseline session was randomized and counterbalanced across the group of ski teams we tested.

The first session after learning group assignment involved a forced exploration. Here, skiers within the learning group were gathered, and the session started by introducing them to the strategies. We explained that we had identified four strategies to enhance racing times on flat slopes in slalom. Subsequently, each strategy was detailed, supported by illustrative drawings in Fig. 1b and corresponding word explanations as outlined in the figure caption and Supplementary Methods C. To confirm comprehension, we conducted two short familiarization trials for each strategy or until the execution met our performance standards. After reviewing the strategies with the skiers, we gathered them in their respective learning group and asked them to rank the strategies (1=best; 4=worst) for what they believe to be the best strategies for improving race times in the flat section of a slalom course. Throughout the instruction and ranking process, skiers were explicitly instructed not to discuss the strategies with each other. It is important to note that the same instructor was used for all learning groups within the tested ski group. After this, the skiers conducted a total of eight trials on the course, with two trials for each strategy. Here, we handled the practice order effect probabilistically by randomizing the order for each skier, under the condition that the first and last four trials tested all strategies. During these trials, the reinforcement learning group received feedback on the timing, whereas the coach provided feedback in the supervised learning groups. After completing the eight rounds, the skiers reevaluated and ranked the strategies.

On the second day, the skiers completed two free choice sessions, each comprising a total of 6 trials in the same slalom courses that were used for the baseline testing the day before. Prior to the 6 free choice trials, the skiers performed one warm-up free skiing run and one warm-up run in the slalom course. In these sessions, the supervised (target skill) learning group consistently selected the theoretically best strategy (that is, ‘extend with rock skis forward’). Conversely, in the supervised (free choice) learning group and the reinforcement learning group, the coach and the skiers, respectively, had the autonomy to choose the skiing strategy for each run. After each session, coaches (except supervised target skills) and skiers were asked to re-evaluate and rank the strategies.

On the third and last day, the skiers performed a retention test and a transfer test to assess the effect of the training approaches on learning and performance. The retention test was performed in the same course as the baseline and acquisition sessions, whereas the transfer test was performed in the transfer course and involved a progression in gate offset from start to finish. Since the transfer test was a new course, we allowed the skiers to inspect the course before the test. The retention and transfer tests were conducted with the three learning groups together. None of the learning groups received any feedback from coaches or time during these tests. After each test, the skiers were asked to rank the strategies.

### 2.5 Analysis

The data were cleaned with custom functions built on tidyverse [53] packages in R [54]. After this process, we validated the data by performing trial counting and visual inspection of the race time to screen for errors. An extensive report of this cleaning and validation process can be found at OSF (https://osf.io/2jxgk).

Due to the hierarchical structure of the data, our general statistical strategy relies on multilevel modeling. At the first level, each skier performed multiple trials during each session. At the second level, each skier was nested within groups of ski teams that performed the experiment together. To account for these multilevel data structures, we leveraged linear mixed-effects models. To model random effects, we adopted a design-driven approach [55, 56], where we sought to account for all nonindependence introduced by repeated sampling from the same ski group and skier. We deployed classical frequentist statistics and fitted these models with the lme4 package [57] in the R [54] programming language. We used a simple coding scheme for our predictors where the intercepts represent the estimated mean of the cell means and the contrasts represent the estimated difference with respect to the reference level, which we set for reinforcement learning. Two-tailed p values and degrees of freedom for each model were derived using the lmerTest package [58] via the Satterthwaite approximation method. Alpha was set to 0.05 for all test statistics.

#### 2.5.1 Race time

Race times were analyzed using linear mixed-effect regression models. Initially, we planned to normalize the racing times by expressing the racing times as the difference from the straight-gliding time performed at the beginning of every session. This difference better approximates the skiers’ actual skill improvement by considering the variance in snow conditions. However, practical considerations led us to deviate from this approach. This change was necessary because we had to flip or shift the course after each day to ensure snow conditions with the least damage. Unfortunately, these adjustments made maintaining a clean, straight-gliding lane difficult since the straight gliding lane crossed many areas with damage to the snow surface (holes) from the previous course set (see Supplementary Methods B for an image of these holes). Collisions with these holes affected the race time, adding noise to the results. Therefore, we used a more conservative approach and analyzed the raw racing times instead of analyzing the normalized racing times.

For the acquisition session, we modeled race time using Session (forced exploration, free choice 1, free choice 2) and Treatment (reinforcement learning, supervised: free choice learning, supervised: target skill learning), and their interactions, as predictors. For retention and transfer, we modeled racing times at these sessions, with treatment added as a predictor. In addition, we used the average performance for each skier on the baseline test as a predictor to improve estimate precision and adjust for any group differences at baseline testing.

To model the effect and development of the strategies we broke the analysis up into different submodels. One analysis focused on differences in the strategies and groups regarding Forced Exploration, where all participants had completed all strategies. Another analysis examined the transition from Forced Exploration to retention for both supervised (free choice) and reinforcement learning. The final model investigated the development between groups specifically for the ”extend with rock skis forward” strategy. Session was coded as a continuous variable in all models.

#### 2.5.2 Strategy choices

Strategy choices were analyzed using generalized linear mixed-effect regression models with a binomial logit-link function. To model the selection of the theoretical best strategy, we inputted the data as logistic, where for each trial (*i*) per skier (*j*) within ski group (*k*), we counted *y_ijk_*= 1 when the skier chose the theoretical best strategy (that is, ‘extend with rock skis forward’) or 0 when they did not. We included Treatment (reinforcement learning, supervised: free choice learning, supervised: target skill learning) and Session (free choice 1, free choice 2, retention, transfer) and their interaction as two variables. To account for the nonindependence of the data structure, we allowed the intercept to vary by including a random intercept for the skier and ski groups.

We adopted the same model formula to model the selection of the estimated best strategy. This time, however, we counted *y_ijk_* = 1 when the skier chose their estimated best strategy and 0 when they did not select that strategy. To estimate the best strategy for each skier in the sample, we used the sample-average method [26] to average the race time for each strategy and selected the strategy with the lowest estimated (that is, best) value. The sessions that we used to form this average were Forced Exploration, Free Choice 1, and Free Choice 2. Due to the scaling issues with generalized linear models, we followed the recommendation to determine the size and significance of the effects of interest using marginal effects on the probability scale [59, 60]. Interactions were assessed using discrete difference (also second difference), which is also in line with these recommendations. To derive these estimates, we used the emmeans package [61].

To learn how the skiers and coaches used feedback to guide their choices, we constructed a ‘win-stay, lose-shift’ model [62–64]. For this analysis, we z-scored the race times for each skier for Free Choices 1 and 2 and counted *y_ijk_* = 1 when the skier repeated the previous strategy and 0 when they did not. The data were modeled using a generalized linear mixed-effect regression model with a binomial logit-link function, with Treatment and z-transformed Race Time and their interaction as the two variables. To test for differences in error sensitivity, we used the marginal effects at the mean derived from the emmeans package [61].

#### 2.5.3 Strategy evaluations and outcomes

To analyze the strategies’ rankings, we used single-level linear regression owing to the singularity of our multilevel models. In this model, we inputted Session as a continuous variable (with the intercept anchored at Familiarization) and Treatment (reinforcement learning, supervised: free choice learning, supervised: target skill learning) as the predictors. For supervised (free choice) learning, we used the coaches’ rankings during the sessions where they selected strategies, and we used the skiers’ rankings when they selected strategies during the retention and transfer tests.

To evaluate how the race times evolved over the next sessions, we had to break down the analysis into three subanalyses because the supervised (target skill) learning group only performed ‘extend with rock skis forward’ during free choice 1 and free choice 2. First, we built at model to assess the difference between the strategies and group differences during forced exploration. To this end, we included Treatment (all learning groups) and Strategy, and their interaction, as factors. For this particular model, we used sliding contrast coding scheme. Second, we built a model to test for differences in improvement on ‘stand against’, ‘rock skis forward’, and ‘extend’ only for the reinforcement and supervised (free choice) learning groups. To this end, we inputted Session as a continuous variable (with the intercept anchored at Familiarization) and Treatment as the predictors. Finally, we built a model to assess the development for all groups on the ”extend with rock skis forward” strategy. This model consisted of Session as a continuous variable (with the intercept anchored at Familiarization) and Treatment as the predictors.

## 3 Results

### 3.1 Race times

We assumed that the strategy choice would greatly impact race time in the slalom course and that the reinforcement learning group would learn to select better strategies and consequently perform better over the course of the experiment compared to the supervised (free choice) learning group where we used the skiers’ own coaches. The supervised (target skill) learning group was instructed to choose the optimal strategy and therefore served as a benchmark for the maximum expected benefit of improved strategy selection through reinforcement learning. As our first step, we analyzed whether the learning groups showed pure time differences across the different sessions without taking the chosen strategy into account.

#### 3.1.1 Greater performance improvement for reinforcement learning compared to supervised (free choice) learning

If the reinforcement learning group learned to select better strategies than the supervised (free choice) learning group, we would expect them to exhibit greater improvement over the course of the three acquisition sessions. Therefore, our hypothesis was that the reinforcement learning group would improve more during the acquisition sessions than the supervised (free choice) learning group. Further, we would expect the supervised (target skill) learning group to rapidly improve race time when the skiers were instructed to only select the theoretical best strategy during the last two acquisition sessions (free choice 1 and free choice 2). We expected that differences in improvement between the learning groups would emerge once the skiers or the coach had the autonomy to pick the strategy over the course of the second (free choice 1) and third (free choice 2) acquisition sessions. Importantly, the three learning groups performed similarly during the forced exploration where they had the same number of trials on each strategy; we found no statistically significant differences between the reinforcement learning group and the supervised (free choice) learning group (*β* = 0.06, 95% CI [-0.2, 0.32], *t*(92.727) = 0.48, *p* = 0.631) or between the supervised (target skill) learning group (*β* = 0.18, 95% CI[-0.08, 0.44], *t*(92.663) = 1.37, *p* = 0.174) during the first acquisition session (forced exploration).

All learning groups significantly improved their race times over the course of the free choice sessions during acquisition (Supplementary Table 1). As expected, the rate at which they improved, differed across the three groups during these two sessions. In line with our expectations, the supervised (target skill) learning group, in which skiers were coached to solely select the theoretically best strategy, showed a statistically significantly greater improvement than the reinforcement learning group from forced exploration to free choice 1 (*β* = -0.12, 95% CI[-0.22, -0.03], *t*(91.777) = -2.58, *p* = 0.012). Conversely, the supervised (free choice) learning group demonstrated a descriptively poorer progression than the reinforcement learning group, albeit the difference was not statistically significant (*β* = 0.08, 95% CI[-0.02, 0.17], *t*(92.5) = 1.61, *p* = 0.110). Continuing this trend, comparing initial performance to performance in the final acquisition session (free choice 2), the reinforcement learning group improved significantly more than the supervised (free choice) learning group did (*β* = 0.14, 95% CI[0.02, 0.26], *t*(95.743) = 2.26, *p* = 0.026). For the same comparison, the supervised (target skill) learning group no longer improved their race times significantly more than the reinforcement learning group (*β* = 0.02, 95% CI[-0.11, 0.14], *t* (95.651) = 0.26, *p* = 0.798). This is due in part to the continued improvement of the reinforcement learning group but also to a descriptive decline from free choice 1 to free choice 2 in the supervised (target skill) learning group, attenuating their initially greater improvement rate. However, we did not find statistical evidence that the reinforcement learning group performed better than the supervised (free choice) or supervised (target skill) learning groups at free choice 1 or free choice 2 (Supplementary Table 2). Fig. 3a presents the mean race time estimates during the three acquisition sessions.

**Fig. 3.**
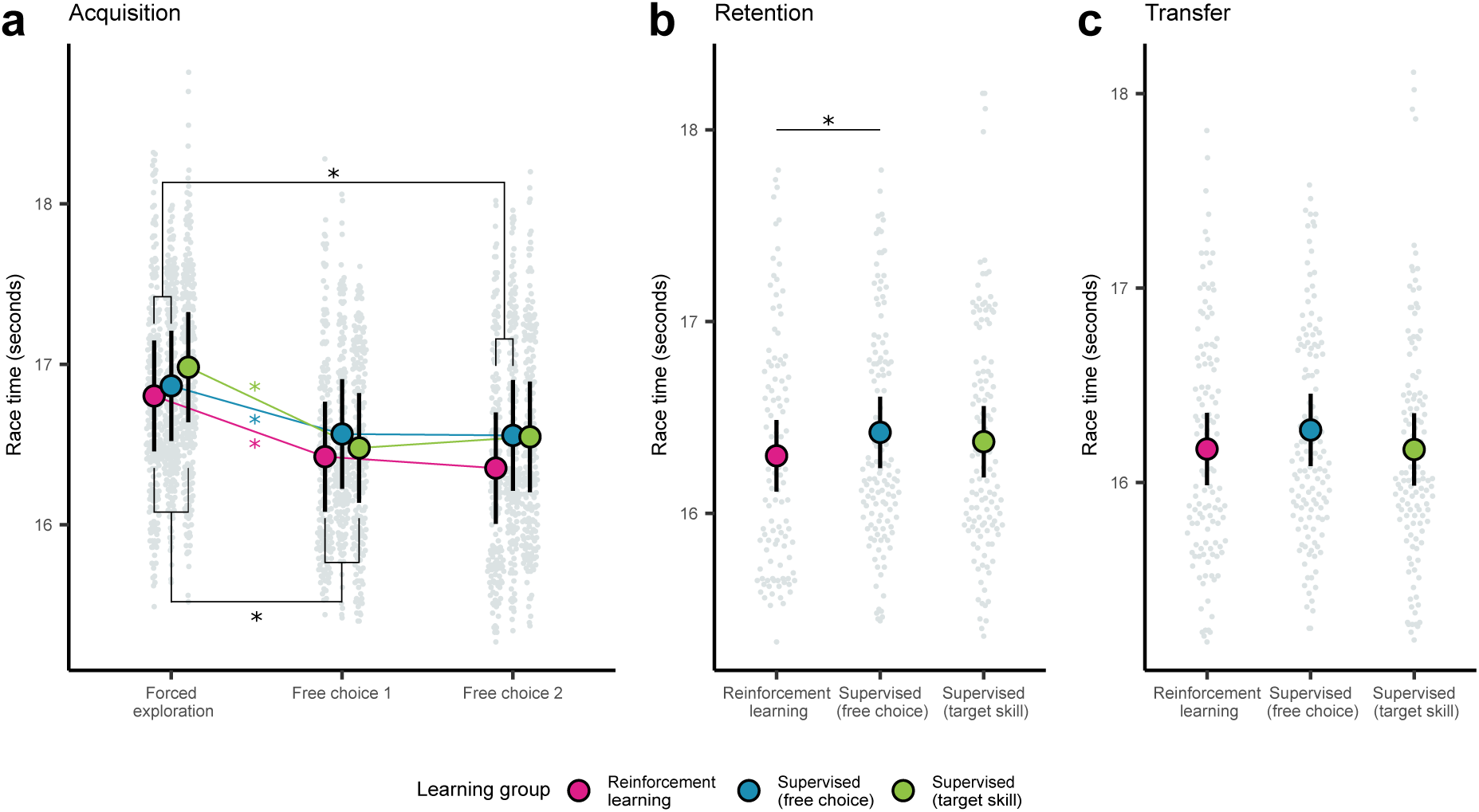
Race time across the different sessions for the three learning groups **a**. Estimated race times during the three acquisition sessions. Forced exploration refers to the sessions wherein skiers tried all strategies, whereas free choice 1 and free choice 2 refer to the session wherein skiers or coaches selected strategies according to their assigned learning groups. **b.** Estimated race times for retention. **c.** Estimated race times for transfer. Intervals represent the 95% confidence intervals (CIs) derived from the models. Asterisks (*) indicate a statistically significant effect. Each light gray point represents a single trial performed by a skier.

#### 3.1.2 Superior retention performance for reinforcement learning compared to supervised (free choice) learning

In the retention session, the skiers independently chose their strategies, irrespective of their assigned groups. We reasoned that at this point, the reinforcement learning group had learned to understand which strategies were most effective and chose them. Therefore, our hypothesis was that the reinforcement learning group would outperform the supervised (free choice) learning group in retention due to better strategy selection learned from observing the race times to evaluate the strategies. Consistent with this hypothesis, we found that the reinforcement learning group performed significantly better than the supervised (free choice) learning group did (*β* = 0.12, 95% CI[0.01, 0.24], *t*(101.422) = 2.12, *p* = 0.037). The difference between the reinforcement learning and supervised (target skill) learning groups also favored reinforcement learning but was smaller and not statistically significant (*β* = 0.07, 95% CI[-0.04, 0.19], *t*(101.63) = 1.27, *p* = 0.206). We therefore provide evidence that the reinforcement learning group performed better at retention than did the supervised (free choice) learning group. Descriptively, the reinforcement learning group performed better, even than the supervised (target skill) learning group, which served as an upper bound reference. Fig. 3b presents the mean race time estimates during retention.

We also hypothesized that reinforcement learning would improve skill transfer to a new slalom course compared to the supervised (free choice) learning group. Similar to retention, the reinforcement learning group performed better than the supervised (free choice) group, yet the difference was smaller and not statistically significant (*β* = 0.1, 95% CI[-0.02, 0.21], *t*(99.979) = 1.7, *p* = 0.091). The race times for reinforcement learning and supervised (target skill) learning groups were on average identical (*β* = 0, 95% CI[-0.12, 0.11], *t*(100.033) = -0.04, *p* = 0.967). Thus, we did not find corroborating evidence for improved transfer. Fig. 3c presents the mean race time estimates during transfer.

### 3.2 Strategy choices

We proposed that the differences in race time between the reinforcement learning group and the supervised (free choice) learning group could be explained by the choice of strategy. Specifically, we hypothesized that the reinforcement learning group would learn to choose better strategies than the supervised (free choice) learning group by learning the strategies’ values directly from observing race times. Fig. 4 displays the percentage selections of the four strategies across all sessions.

**Fig. 4.**
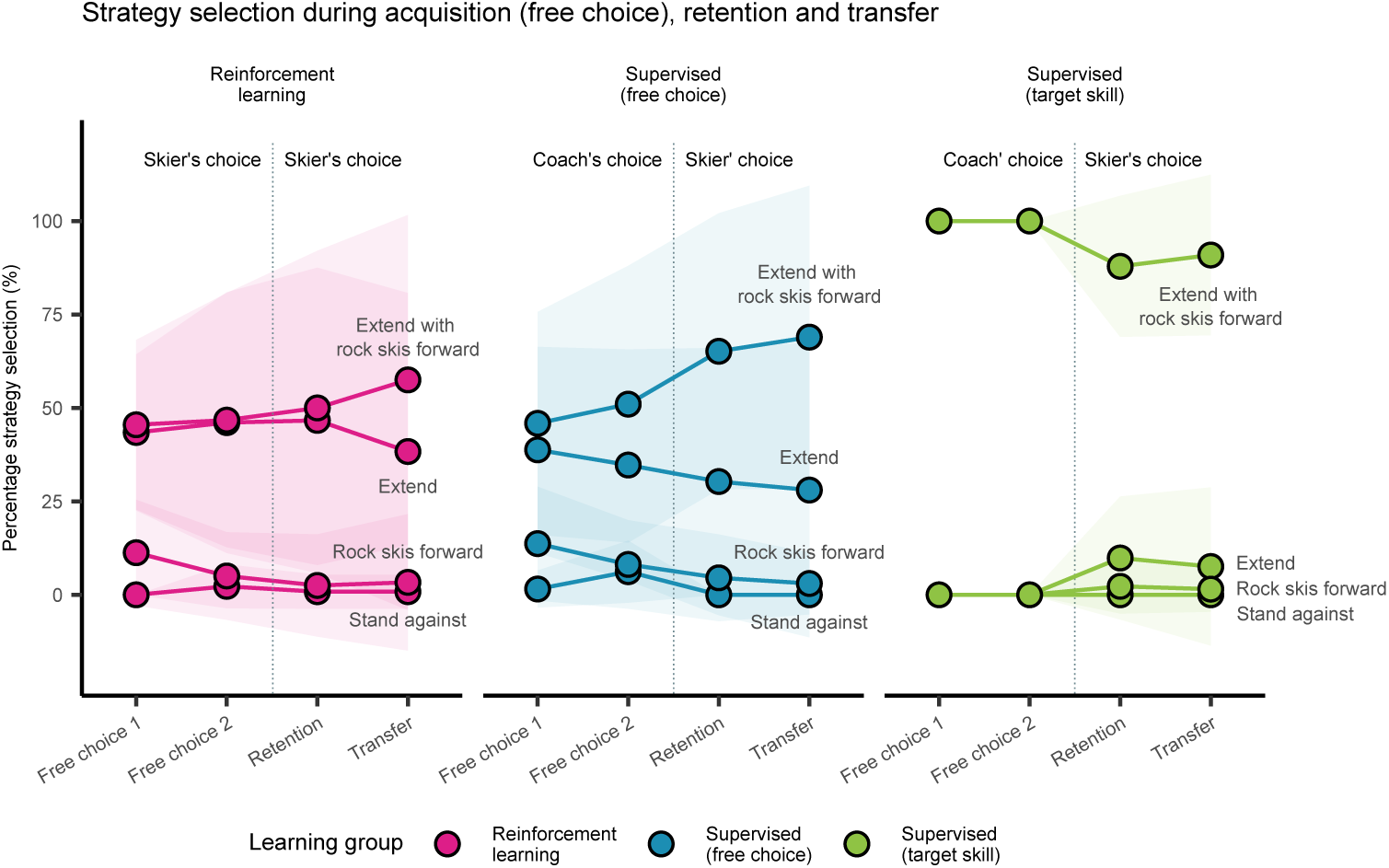
Percentage of choices for each strategy and session (circles). Shaded error bars represent standard deviations.

#### 3.2.1 No greater increase in selecting the theoretical best strategy for reinforcement compared to supervised (free choice) learning

We assumed that the skiers performed best with the strategy we defined as the theoretically best strategy (that is, with the ”extend with rock skis forward”) and tested whether the reinforcement learning group developed a greater probability of selecting this strategy than the supervised (free choice) learning group. We found that both learning groups showed a statistically significant increase in the probability of choosing the strategy we considered theoretically optimal across the four sessions (Supplementary Table 3); however, the timing of this increase differed between the two groups. For the supervised (free choice) learning group we found a statistically significant increase from free choice 1 to retention, when skiers in this group were given autonomy to choose strategies themselves (0.31, 95% CI[0.18, 0.44], *z* = 4.59, *p <* 0.001). This probability increase was significantly greater than the increase for the reinforcement learning group (0.24, 95% CI[0.05, 0.43], *z* = 2.43, *p* = 0.015), which did not significantly increase from free choice 1 (0.07, 95% CI[-0.07, 0.21], *z* = 0.96, *p* = 0.339). The reinforcement learning group, on the other hand, significantly increased the probability of choosing the theoretically best strategy from free choice 1 to the transfer session(0.18, 95%CI[0.04, 0.32], *z* = 2.58, *p* = 0.010). Despite the descriptively higher probability of choosing the theoretically best strategy in the supervised (free choice) learning group during the free choice 2, retention and transfer sessions, none of the differences between the groups at each session were statistically significant (Supplementary Table 4). Fig. 5a displays the predicted probabilities for the learning groups across the four sessions where the skiers or coaches were given autonomy to select strategies themselves. Note that the supervised (target skill) learning group was excluded from this analysis because by definition only the theoretically best strategy was selected.

**Fig. 5.**
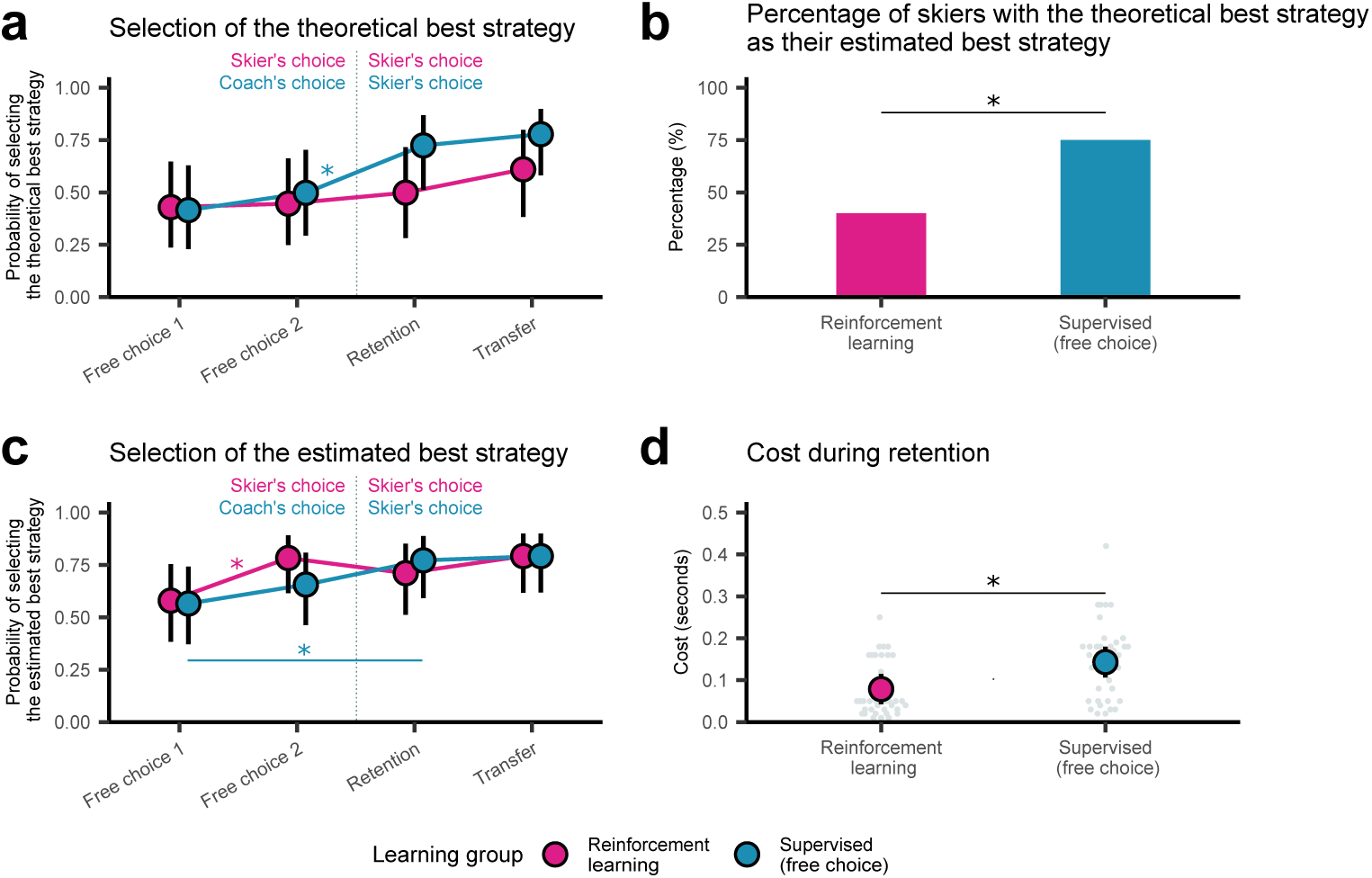
Strategy selection for reinforcement learning (pink) and supervised (free choice) learning (blue) groups during acquisition. **a.** Predicted probability of choosing the theoretically best strategy (that is, ‘extend with rock skis forward’). **b.** Percentage of skiers performing best using the theoretically best strategy. **c.** Predicted probability of selecting the individual skier’s estimated best strategy. **d.** Estimated cost of selecting a suboptimal strategy during retention. **a., c.** Intervals represent the 95% confidence intervals (CIs) derived from the models. Asterisks (*) indicate a statistically significant effect.

#### 3.2.2 Differences in strategy performance between reinforcement and supervised (free choice) learning groups

Based on the previous analysis, the reinforcement learning group did not develop a greater probability of selecting what we defined as the theoretically optimal strategy compared to the supervised (free choice) learning group. Instead, the descriptive trend favored supervised (free choice) learning. Therefore, we wondered whether a greater proportion of skiers in the supervised (free choice) learning group had the theoretical best strategy as their estimated best strategy, and performed a followup analysis to test this possibility. Overall, 75% of the skiers in the supervised (free choice) learning group performed best using the theoretically best strategy compared to only 40% in the reinforcement learning group. A chisquare test revealed a statistically significant difference between groups *χ*^2^ = 6.42, *p* = 0.01). Fig. 5b shows the proportion of skiers for each group that performed best with the theoretical best strategy. Note that skiers in the supervised (target skill) group only used the theoretically best strategy, so they could not be included in this analysis. The observation that more skiers in the supervised (free choice) learning group had the theoretically best strategy as their estimated best strategy could explain why they had a descriptively greater probability of choosing it compared to the reinforcement learning group.

#### 3.2.3 No greater increase in selecting individual skier’s estimated best strategy for reinforcement compared to the supervised (free choice) learning

We did not find evidence that the reinforcement learning group developed a greater predicted probability of choosing the theoretically optimal strategy. However, this strategy was not the best strategy for every skier. Therefore, what if we instead base the analysis on the individual skier estimated best strategy? Would the reinforcement learning group have a higher predicted probability of choosing this strategy? During the first session, in which we allowed skiers and coaches to choose their own strategies (free choice 1), we found no statistically significant differences between the groups (0.01, 95% CI[-0.22, 0.24], *z* = 0.12, *p* = 0.904). Both groups significantly improved their choices over the course of the sessions relative to free choice 1 (Supplementary Table 5), but the significant improvement came at different time points. We found a statistically significant improvement in choice for the reinforcement learning group from free choice 1 to free choice 2 (0.2, 95% CI[0.09, 0.32], *z* = 3.39, *p <* 0.001), but not for the supervised (free choice) learning group (0.09, 95% CI[-0.03, 0.21], *z* = 1.48, *p* = 0.140). However, the reinforcement learning group did not increase their probability of choosing their individually best strategy significantly more than the supervised (free choice) group (0.11, 95% CI [-0.28, 0.05], *z* = -1.33, *p* = 0.184). For the supervised (free choice) learning group we found a statistically significant improvement in their choices when the skiers made their own strategy choices during retention (0.21, 95% CI[0.08, 0.34], *z* = 3.11, *p* = 0.002). We found no statistically significant differences between the groups in any of the sessions (Supplementary Table 6). Therefore, we did not find corroborating evidence that the reinforcement learning group learned to pick the best strategy more often than the supervised (free choice) learning group. Fig. 5c displays the predicted probabilities of choosing the estimated best strategy for the learning groups across the sessions.

#### 3.2.4 Lower cost of choosing suboptimal strategies for reinforcement compared to supervised (free choice) learning during retention

Selecting an effective movement strategy can extend beyond the mere selection of the single best strategy, however. Sometimes, several strategies produce outcomes that are nearly identical, giving athletes the flexibility to select any option without sacrificing performance. The task then becomes one of selecting strategies that offer comparable outcomes while avoiding those that carry the risk of substantially worse performance. We reasoned that discerning differences between fairly similar strategies could be tricky for skiers due to small time differentials that are influenced by noise, such as hitting a bump. Consequently, two or more strategies may appear nearly identical and be difficult for skiers to distinguish. We proposed that the reinforcement learning group had learned to select good strategies that all led to similarly good outcomes and learned to steer away from those that led to poor outcomes. To test this idea, we calculated the expected difference between a skier’s chosen ‘suboptimal’ strategy and their estimated best strategy, which we termed ‘cost.’ Using this measure, we ran a follow-up analysis to test whether the reinforcement learning group had a lower ‘cost’ during retention than did the supervised (free choice) learning group. We limited the testing to retention, as it was only in this session that the skiers in the supervised (free choice) learning group were allowed to choose their own strategy and who were in the same slalom course they had previously skied. This analysis revealed that the reinforcement learning group had significantly lower costs of selecting suboptimal strategies during retention than the supervised (free choice) learning group (*β* = 0.06, 95% CI [0.01, 0.12], *t*(30.789) = 2.55, *p* = 0.016). This result suggests that skiers in the reinforcement learning group may have learned to select better strategies, although we did not find evidence that they had a greater probability of picking the best strategy. Fig. 5d displays the expected cost for the suboptimally chosen strategies.

#### 3.2.5 Similar use of feedback for strategy selection in reinforcement and supervised (free choice) learning groups

Finally we examined the extent to which decision-makers (skiers in reinforcement learning and coaches in supervised learning) engaged with their feedback times to inform decisions about which strategy to select next. Our hypothesis was that, compared with coaches, skiers in the reinforcement learning group would engage more actively with these race times to improve their choices. To test this hypothesis, we conducted a ‘winstay, lose-shift’ analysis, where the primary assumption is that all information used for decision-making stems from the last trial (n-1)[63, 64]. In this analysis, heightened sensitivity to the most recent feedback is reflected by a high predicted probability of repeating an action following positive feedback (a fast time, compared to the average time) and a low predicted probability following negative feedback (a slow time, compared to the average time) on the preceding trial. We found statistically significant estimated marginal effects at the mean for both the reinforcement learning group (-0.18, 95% CI[-0.26, -0.11], *z* = -4.8, *p <* 0.001) and the supervised (free choice) learning group (-0.11, 95% CI[-0.17, -0.04], *z* = -3.29, *p <* 0.001). These findings suggest that both groups were more likely to repeat a strategy if the previous trial feedback was good. Despite the large descriptive difference in the marginal effect between groups, this difference was not statistically significant (-0.08, 95% CI [-0.17, 0.02], *z* = -1.55, *p* = 0.121). Thus, although we found that both learning groups were sensitive to the most recent feedback (either using it themselves to determine choices, or having the coach use it to determine choices), we did not find evidence that the reinforcement learning group had greater sensitivity than the supervised (free choice) learning group. Fig. 6 shows the predicted probability of repeating the strategy on the previous trial depending on the feedback.

**Fig. 6.**
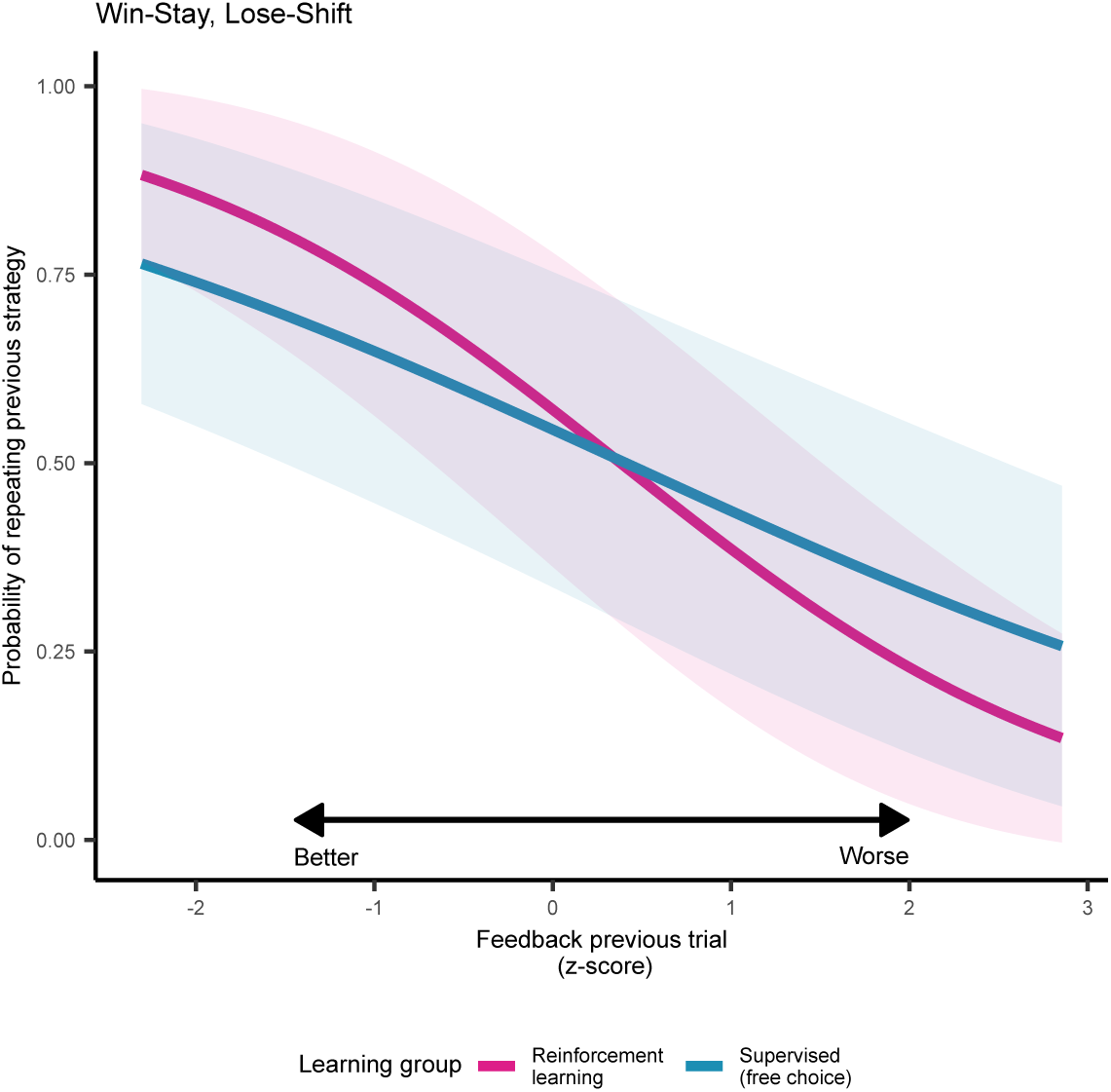
Win-stay, lose-shift comparison between reinforcement learning and supervised (free choice) learning. The line shows the predicted probability of repeating the previously chosen strategy based on its trial feedback, along with a 95% CI in the shaded error bar

### 3.3 Strategy evaluations and outcomes

As a final step, we assessed how the learning groups’ understanding of the strategies evolved throughout the experiment and how well this understanding aligned with the actual times achieved for the four strategies. The purpose of this analysis was to learn how the learning groups’ understanding of the strategies governed their choices. To assess this, we divided the analysis into two parts. First, we assessed the learning group’s evaluation of the strategies. Second, we assessed the effect of the learning group’s race on the participants’ strategies.

#### 3.3.1 Greater separation in strategy evaluations for reinforcement compared to supervised (target skill) learning

To assess how the learning groups’ knowledge of the strategies evolved and governed their choices we asked the skiers and coaches (excluding coaches involved in supervised (target skill) learning) to evaluate the strategies by ranking them from best (1) to worst (4). They first ranked the strategies upon their introduction (familiarization) to the strategies and then after each session. We expected that the reinforcement learning group would develop more distinct evaluations of the strategies compared to the supervised learning groups by having the race times at their disposal to evaluate the strategies. To evaluate this, we gathered the coaches’ rankings in the supervised (free choice) learning group for analysis during the two acquisition sessions, as they were responsible for selecting the strategy during these sessions. Conversely, we collected rankings from the skiers in the supervised (free choice) learning group for analysis during the retention and transfer sessions, as they were responsible for making decisions in these sessions. For supervised (target skill) learning, we exclusively focused on the skiers’ rankings since the coach had been influenced by the information we gave them about the strategies before the experiment.

After the introduction to the strategies, but before the skiers had the chance to properly try out the strategies on the slalom course (familiarization), we found that all groups ranked ‘extend with rock skis forward’ as the best, followed by ‘extend,’ ‘rock skis forward,’ and ‘stand against’ (Supplementary Table 7 and Fig. 7a). We did not find any statistically significant difference between groups during this familiarization (Supplementary Table 8), with two exceptions: the supervised (target skill) learning group ranked ‘extend’ worse (*β* = 0.45, 95% CI[0.24, 0.67], *t*(1700) = 4.17, *p <* 0.001) and ‘extend with rock skis forward’ better (*β* = -0.42, 95% CI[-0.63, -0.2], *t*(1700) = -3.83, *p <* 0.001) than the reinforcement learning group.

**Fig. 7.**
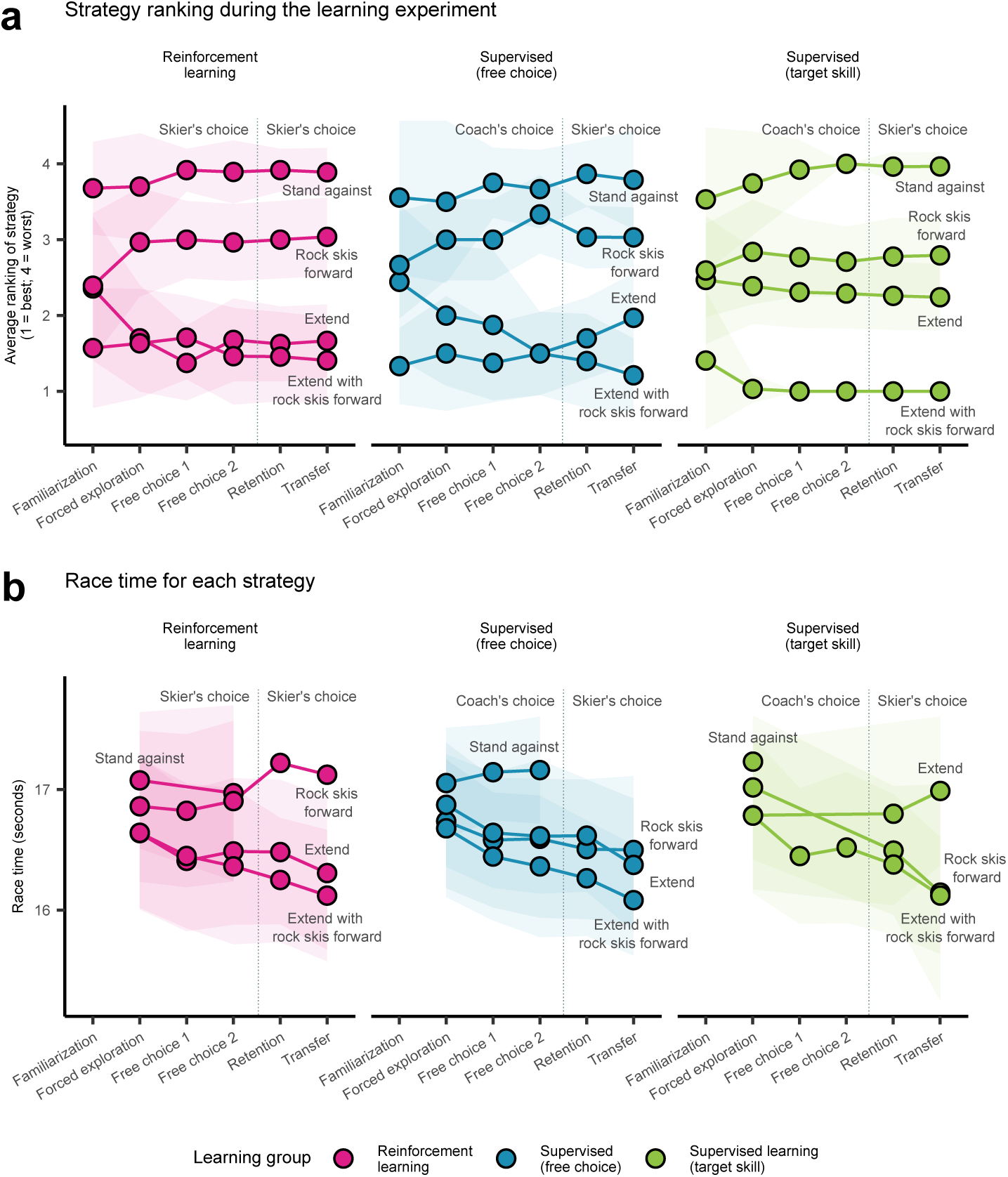
Strategy evaluations and performance. **a.** Average descriptive ranking of the four strategies per treatment group. Rankings range from 1 (best) to 4 (worst). For supervised (free choice) learning, the coach’s ranking during the acquisition phase and the skier’s ranking during the retention and transfer phases are plotted, reflecting the decision-maker for strategy selection. The circle represents the mean, and the shaded area indicates the standard deviation (SD). **b.** Average race time of the four strategies across the three learning groups. The circle represents the mean, and the shaded area represents the SD. Note that all skiers tested the strategies during the forced exploration phase, but as the study progressed, there may have been fewer observations for some strategies. Consequently, the calculation of the mean might be heavily influenced by these observations.

Over the course of the sessions, we observed notable shifts in the average rankings of the strategies (Supplementary Table 9). The change was relatively flat and unchanged in terms of position for the worst (’stand against’) and best (’extend with rock skis forward’) strategies. However, we found that ‘stand against’ was ranked significantly worse (*β* = 0.09, 95% CI[0.04, 0.13], *t*(1700) = 3.4, *p <* 0.001) and that ‘extend with rock skis forward’ was ranked significantly better (*β* = -0.06, 95% CI[-0.11, -0.01], *t*(1700) = -2.51, *p* = 0.012) over time in the supervised (target skill) learning group. Although the reinforcement learning and supervised (free choice) learning groups followed the same trend, these trends were not statistically significant. More marked shifts were observed for the two middle-ranked strategies: ‘extend’ and ‘rock skis forward’. Specifically, the reinforcement learning group ranked ‘extend’ significantly better (*β* = -0.1, 95% CI[-0.15, -0.05], *t*(1700) = -3.73, *p <* 0.001) and ‘rock skis forward’ significantly worse (*β* = 0.09, 95% CI[0.04, 0.15], *t*(1700) = 3.57, *p <* 0.001) over the course of the sessions. The supervised (target skill) and supervised (free choice) groups also had the same trend, but the magnitude was smaller and did not reach statistical significance (Supplementary Table 9). Interestingly, we found that there was a greater change in the ranking of ‘rock skis forward’ in the reinforcement learning group than in supervised (target skill) learning group (*β* = -0.07, 95% CI[-0.14 to 0], *t*(1700) = -1.97, *p* = 0.049), suggesting a larger shift in strategy ranking for this strategy in the reinforcement learning group. Overall, these results show that the groups updated their evaluations of the different strategies based on their experience and that the reinforcement learning group did so more than the supervised (free choice) learning group.

#### 3.3.2 Strategy performance mirrors strategy evaluation

We would expect that, at least toward the later sessions, these evaluations reflect the race times for each strategy. To determine whether the pattern in strategy ranking aligned with race time for the strategies, we tested how performance changed across sessions for each strategy. Overall, race times for each strategy mirrored each group’s strategy evaluations (Fig. 7b). Note that skiers’ evaluations are also reflected in the fact that there are no race times for the worst-rated strategy during retention and transfer in any of the groups, because skiers did not select this strategy. During the first acquisition session when skiers tested all strategies (forced exploration), we found that all groups on average performed descriptively better with ‘extend with rock skis forward’, followed by ‘extend’, ‘rock skis forward’, and finally ‘stand against’ (Supplementary Table 10). This finding largely aligns with the rankings given for each strategy during this session (see section 3.3.1).

We found no statistically significant differences in performance between the reinforcement learning and supervised learning groups on any of the strategies (Supplementary Table 11). To evaluate how the race times evolved over the next sessions, we had to break down the analysis into two subanalyses because the supervised (target skill) learning group only performed ‘extend with rock skis forward’ during free choice 1 and free choice 2. First, we assessed the development for all groups on the ”extend with rock skis forward” strategy. This analysis revealed that all groups improved on the ”extend with rock skis forward” strategy over the course of the sessions (Supplementary Table 12). However, we found no statistically significant differences in skill improvement for this strategy between the reinforcement learning group and either supervised (free choice) learning group (*β* = 0, 95% CI[-0.03, 0.02], *t*(1064.204) = - 0.3, *p* = 0.765) or supervised (target skill) learning group (*β* = 0.02, 95% CI [-0.01, 0.04], *t*(1062.577) = 1.07, *p* = 0.283). This result shows that performing all trials with the optimal strategy did not further improve the performance of the supervised (target skill) learning group. Second, we tested for differences in improvement on ‘stand against’, ‘rock skis forward’, and ‘extend’ only for the reinforcement and supervised (free choice) learning groups. We found a statistically significant improvement in all strategies over the course of the sessions to retention, except for ‘stand against’ in the reinforcement learning group (Supplementary Table 13). This deviation, however, may be attributed to the limited number of observations for this strategy within this group.

Interestingly, we found that the reinforcement learning group improved more on the ”extend” strategy than did the supervised (free choice) learning group (*β* = 0.04, 95% CI[0.01, 0.07], *t*(1378.879) = 2.34, *p* = 0.020). This result aligns well with the reinforcement learning group’s evaluation of this strategy, which they ranked significantly better over the course of the sessions, whereas the supervised (free choice) learning group did not(see above); and this result might explain the reduced cost of selecting a suboptimal strategy. Notably, the reinforcement learning group was not assigned a coach to help them improve on that strategy, but could only rely on their race times. We did not find evidence for any interaction effect for the other two strategies (Supplementary Table 14). Taking the evaluations and outcomes collectively, it seems that these evaluations reflected the race times well for each strategy.

## 4 Discussion

Skilled athletes need powerful movement strategies to solve tasks effectively. Typically, athletes learn these strategies with instruction-based teaching methods where coaches offer athletes a correct solution. Inspired by insights from decision neuroscience, we asked whether skilled athletes can instead improve strategy decisions and consequently enhance performance by shifting from an instruction (supervised learning) to an evaluation-oriented (reinforcement learning) training strategy. In a three-day skill learning experiment, we found that the reinforcement learning group achieved greater improvement over the course of the acquisition sessions and performed better during the retention session than did the supervised (free choice) learning group, where the coach selected a strategy for the skier. Interestingly, the reinforcement learning group also performed descriptively better than the supervised (target skill) learning group, which was meant to serve as a benchmark for the upper limit of performance achievable through optimal strategy choices. However, we found less convincing evidence corroborating our hypothesis that reinforcement learning improved transfer.

At a broad level, our findings dovetail with two research streams that have shown that withholding instruction or feedback can be an effective way to enhance skill learning. First, our finding that the reinforcement learning group showed better skill performance during retention than the supervised (free choice) learning group echoes earlier results from ‘discovery learning’ approaches showing that instructional methods do not always benefit learning motor skills [24, 25, 65, 66]. Second, this finding mirrors previous studies that have found better skill retention from reinforcement learning-based interventions [35, 37, 38, 67]. We expand these two lines of research to a complex sport involving skilled athletes and show that teaching strategies other than instruction can be effective for skill learning.

We proposed that the reinforcement learning group’s improved performance compared to the supervised (free choice) learning group resulted from better strategy choices. However, we found no corroborating evidence for this hypothesis when better strategy choices were defined as choices of a single best strategy; the reinforcement learning group did not begin with or develop a greater probability of choosing the theoretically optimal strategy or the individual skier’s estimated best strategy compared with the supervised (free choice) learning group. Instead, we found that the skiers in the reinforcement learning group who made ‘suboptimal’ choices during the retention session incurred a lower ‘cost’ for their choices than did the skiers in the supervised (free choice) learning group. The reinforcement learning group therefore appears to have learned to select strategies that offered comparable outcomes and to avoid those that carried the risk of substantially worse performance. One possibility is that the reinforcement learning group discovered that the ”extend” strategy alone almost provided full benefits on its own and chose this strategy because it was simpler than ”extend with rock skis forward”.

Improved strategy selection appears not to be the sole explanation for the improved performance of the reinforcement learning group, however. Two other key findings were that the reinforcement learning group had a lower proportion of skiers who performed best with ‘extend with rock skis forward’ than did the supervised (free choice) learning group, and that the reinforcement learning group improved more on the ‘extend’ strategy over the course of the sessions than did the supervised (free choice) learning group. In conjunction these findings suggest that the reinforcement learning group developed the ‘extend’ strategy in an unanticipated direction based on the race time feedback. Comments from a few coaches, who watched the retention and transfer from the sideline, mentioned that skiers in the reinforcement learning group used more forceful arm movements than did the skiers in the other groups, although the instructions did not explicitly tell them to do that. As such, it seems that the reinforcement learning group refined the ‘extend’ strategy to a new level. Unfortunately, we were not able to precisely quantify how this ‘new’ strategy was executed with our measurements.

What information guided the strategy decisions in the two learning groups? We found that both learning groups’ decisions could be accounted for by a ‘winstay, lose-shift’ heuristic, where the decision maker had a greater probability of continuing with a chosen strategy if the performance was better than the skier’s average and conversely was more likely to switch strategies following negative feedback. Such ‘win-stay, lose-shift’ patterns have also been found in a previous study on skill learning [68] and appear to be a general characteristic of learning to make good decisions to achieve goals. Note that it is possible that learners used more that just the most recent feedback to inform strategy selection, as would be expected in reinforcement learning. Differentiating between ‘win-stay, lose-shift’ and more complex learning and decision-making strategies is beyond the scope of the current manuscript. Although the reinforcement learning group showed increased sensitivity to the most recent feedback compared to the supervised (free choice) group, this difference was not statistically significant. A potential reason for this is that the coaches in the supervised (free choice) learning group were highly experienced coaches with a great deal of knowledge, and they also had access to the time. Future studies might consider investigating different coach groups and with and without access to times to improve the understanding of how these decisions are made.

Contrary to our expectation, we did not find convincing evidence that reinforcement learning improved transfer compared with either of the two supervised learning groups. One explanation for this could be that reinforcement learning only improves performance in situations where rewards have been previously received [69], and aligns with [38] who found that reinforcement learning improved retention but not transfer compared with supervised learning. However, we can not argue for differential effects on retention and transfer in our data. The estimated mean difference between the reinforcement learning group and the supervised (free choice) learning group was quite similar to that during the retention session (0.12 sec. on retention versus 0.1 sec. on transfer), and the p value was also low (0.091). By definition, athletes are expected to show greater variability when transitioning from retention to transfer testing because the athlete are exposed to a new situation. Therefore, it is possible that our sample does not include a sufficient number of skiers to achieve adequate statistical power to detect such an effect. Alternatively, a more varied learning approach, where learners are exposed to frequent switches between situations (in our case, slalom courses), may be necessary to grasp the task’s structure and promote transfer [70, 71]. Future research should address the link between reinforcement learning and transfer more closely.

Interestingly, despite the broad consensus that ‘extend with rock skis forward’ was the best strategy upon the introduction of these strategies, coaches and skiers faced greater challenges in determining which of the two strategies, ‘rock skis forward’ and ‘extend’, was the second best. Over the course of the next sessions, we found that the reinforcement learning group learned to distinguish these strategies from each other, as evidenced by a significantly improved assessment of ‘extend’ and a poorer assessment of ‘rock skis forward’, which we did not find in the other two learning groups. In addition, we found that the reinforcement learning group ranked “rock skis forward” more poorly over the course of the sessions than did the supervised (target skill) learning group. Learners may miss potential learning by only being exposed to one strategy instead of a broader exploration of alternatives [72–74], as was the case with the supervised (target skill) learning group. Being exposed to reinforcement learning-type training may stimulate reasoning processes in learners due to task variability that enables them to cultivate a deeper comprehension of the relationship between their actions and performance outcomes. Over time, improving such reasoning capacity could prove crucial for developing innovative and truly brilliant solutions to achieve task goals [14]. It is possible that reinforcement learning, when used to train learners to choose between two or more explicit strategies, can promote reasoning processes that may be important for developing expertise. Future studies should test this idea. Our findings have important implications for coaches when designing training sessions to enhance skills. We advise coaches to formulate explicit strategies to enhance performance in their sport and allow learners to directly learn the action values of these strategies by trying them out and comparing them. Doing so could help learners better understand the functions and effects of various strategies and enable them to make better strategy choices. However, we do not propose replacing traditional teaching methods with this approach but suggest integrating it as a supplementary tool to improve strategic decisions and enhance athletes’ understanding of how their actions influence outcomes.

Before practitioners embrace our recommendation to incorporate more strategy evaluations into their coaching or teaching practices, it is important to consider the practical significance of the effect size and its potential amplifying and counteracting mechanisms[75]. Notably, the estimated effect size during the retention session was smaller than our predefined smallest effect size of interest. This benchmark, however, was set for a longer slalom course (that is, more gates) and more training sessions than we could execute due to space and time constraints in the ski hall. Consequently, we exercise caution in outright dismissing its practical significance. Instead, if we assume that a full slalom course of approximately 45 seconds has a flat section equivalent to approximately 15 seconds (which is quite typical) and that a slalom race consists of two runs, the 0.12-second difference can translate into an improved Fédération Internationale de Ski (FIS) world ranking of 27 positions for females and 65 for males, based on a median ranking of 600 in our sample (see Supplementary Discussion). Practitioners could therefore achieve meaningful gains by applying reinforcement learning to train strategic decisions. However, we must remember that sporting situations are diverse and erratic, often requiring a athlete to switch between different strategies during a performance. The ability to successfully implement these adaptive switches is a hallmark of sporting expertise [1, 6, 8]. Our study did not capture such decisions because we focused only on flat slopes. Finally, it is also possible that the estimated group mean differences could have been greater had the reinforcement learning group and the supervised learning groups undergone retention and transfer sessions separately, thereby preventing any information leakage. With our current procedure, it is conceivable that the skiers observed each other, potentially leading to treatment diffusion. However, this decision was made to mirror the conditions of alpine competitions and gave us confidence that athletes experienced similar conditions during testing.

### 4.1 Limitation of the study

We did not have motion capture data available to analyze the skiers’ execution of the strategies. Therefore, we cannot comment on the actual execution of the strategies, other than verifying that they were able to perform the strategies during the familiarization test. However, taking into account execution might have led to explosion of actually implemented strategies that would then be difficult to study. Indeed, considering a small set of discrete strategies, as we did in this study, has been the recommended approach for studying action selection for motor skill learning [9, 76]. Another limitation is the relatively short duration of the learning experiment. The idea, however, was to mimic a typical short training session for alpine skiers and study whether we could find meaningful effects within this short time window.

## 5 Conclusion

In summary, our data showed better learning with reinforcement learning than with the traditional instruction-based coaching approach, but the advantage did not transfer to a new slalom course. Only by informing coaches about what we believed to be the best strategy for improving race time on flat sections in slalom (based on mechanics and evidence from the field) were we able to achieve learning effects comparable to those of reinforcement learning. Interesting, although the supervised (target skill) learning group always picked the correct strategy, they did not improve more on this strategy than the reinforcement learning group. In accordance with previous suggestions [77, 78], our findings suggest that reinforcement learning can be an important learning strategy for improving strategy choices and learning in skilled athletes.

## Supplementary information

## Acknowledgements

We thank SNØ (https://snooslo.no/) and Igloo Innovation (https://iglooinnovation.no/) for allowing us to use the indoor ski hall and facilities for testing. We thank Christian Mitter for helping us define the strategies and giving us ideas for how to communicate them to the skiers. We thank Erland Hoff Thommasen, Jeremy Phyffer, Mai-Sissel Linløkken, Erland Vedeler Stubbe, Johan Ansnes, Ørjan Lydersen, William Farstad Olsen, Victoria S. Placht, Einar Witteveen, Synne Sofie Stangeland, Kasper Sjøstrand, Tom Knudsen, and Tim Gfeller for helping with watering the hill and assisting during the data collection. We thank Tine SA for providing the skiers with food and nutrients during the data collection. We thank all the ski coaches and skiers who participated and made this study possible. We also want to thank Andrea Pisauro, Doron Cohen, Jørgen Jensen, and Christian Thue Bjørndal for helpful comments and feedback on our preprint. The first author would also like to give special thanks to Cameron Patrick, Isabella R. Ghement and James Steele for their general methodological and statistical discussions.

## Declarations

### 5.1 Data and code availability

The datasets generated during and/or analysed during the current study are available in the Open Science Framework repository repository, https://osf.io/uw986/.

### 5.2 Author contribution

C.M. led conception and design, led the data collection, analyzed and interpreted the data, drafted and revised the article, and agreed to the submitted version for publication. M. G., S. L.T.,T.L ., P.H. and R.R. contributed to the design, data collection, and revised and agreed to the submitted version for publication. R. F. supervised C.M. in the conception and design, analyzed and interpreted the data, revised the article, and agreed to the submitted version for publication.

