## Supplemental information for "Reinforcement learning enhances training efficiency in high-performance athletes"

### Supplementary Material

#### Supplementary Table

**Supplementary Table 1**: Race time: Estimated improvement across sessions in acquisition

| Term | Estimate | SE | df | CI | t | p |
|---|---|---|---|---|---|---|
| (Intercept) | 16.62 | 0.13 | 3.93 | 16.27-16.97 | 132.69 | < 0.001 |
| supervised (free choice) | 0.14 | 0.13 | 85.83 | -0.12-0.39 | 1.04 | 0.299 |
| supervised (target skill) | 0.14 | 0.13 | 85.82 | -0.12-0.40 | 1.10 | 0.276 |
| reinforcement learning : free choice 1 | -0.38 | 0.03 | 91.63 | -0.45–0.31 | -10.82 | < 0.001 |
| supervised (free choice) : free choice 1 | -0.30 | 0.03 | 93.47 | -0.37–0.23 | -8.96 | < 0.001 |
| supervised (target skill) : free choice 1 | -0.50 | 0.03 | 91.95 | -0.57–0.44 | -15.08 | < 0.001 |
| reinforcement learning : free choice 2 | -0.45 | 0.05 | 95.16 | -0.54–0.36 | -9.99 | < 0.001 |
| supervised (free choice) : free choice 2 | -0.31 | 0.04 | 96.39 | -0.39–0.22 | -7.17 | < 0.001 |
| supervised (target skill) : free choice 2 | -0.43 | 0.04 | 96.20 | -0.52–0.35 | -10.07 | < 0.001 |
| sd(Intercept) | 0.51 | | | | | |
| cor(Intercept,free choice 1) | -0.03 | | | | | |
| cor(Intercept,free choice 2) | 0.23 | | | | | |
| sd(free choice 1) | 0.15 | | | | | |
| cor(free choice 1, free choice 2) | 0.73 | | | | | |
| sd(free choice 2) | 0.22 | | | | | |
| sd(Intercept) | 0.23 | | | | | |
| sd(Observation) | 0.21 | | | | | |

Formula: racetime $\sim$ treatment / session + (1 |skigroup) + (1 + session | skigroup:skier)

**Supplementary Table 2**: Race time: Estimated difference between groups during acquisition

| Term | Estimate | SE | df | CI | t | p |
|---|---|---|---|---|---|---|
| (Intercept) | 16.62 | 0.13 | 3.93 | 16.27-16.97 | 132.68 | < 0.001 |
| free choice 1 | -0.39 | 0.02 | 92.32 | -0.43–0.35 | -20.09 | < 0.001 |
| free choice 2 | -0.40 | 0.03 | 95.89 | -0.45–0.35 | -15.74 | < 0.001 |
| forced exploration : supervised (free choice) | 0.06 | 0.13 | 92.73 | -0.20-0.32 | 0.48 | 0.631 |
| free choice 1 : supervised (free choice) | 0.14 | 0.13 | 86.15 | -0.11-0.39 | 1.11 | 0.272 |
| free choice 2 : supervised (free choice) | 0.20 | 0.14 | 78.00 | -0.08-0.49 | 1.43 | 0.157 |
| forced exploration : supervised (target skill) | 0.18 | 0.13 | 92.66 | -0.08-0.44 | 1.37 | 0.174 |
| free choice 1 : supervised (target skill) | 0.05 | 0.13 | 86.12 | -0.20-0.31 | 0.43 | 0.672 |
| free choice 2 : supervised (target skill) | 0.19 | 0.14 | 78.02 | -0.09-0.48 | 1.37 | 0.176 |
| sd(Intercept) | 0.51 | | | | | |
| cor(Intercept, free choice 1) | -0.03 | | | | | |
| cor(Intercept, free choice 2) | 0.23 | | | | | |
| sd(free choice 1) | 0.15 | | | | | |
| cor(free choice 1, free choice 2) | 0.73 | | | | | |
| sd(free choice 2) | 0.22 | | | | | |
| sd(Intercept) | 0.23 | | | | | |
| sd(Observation) | 0.21 | | | | | |

Formula: racetime ∼ session / treatment + (1 |skigroup) + (1 + session | skigroup:skier)

**Supplementary Table 3**: Strategy choice: Probability change across sessions

| Contrast | Treatment | Estimate | SE | df | CI | z | p |
|---|---|---|---|---|---|---|---|
| free choice 2 - free choice 1 | rl | 0.02 | 0.06 | Inf | -0.11-0.14 | 0.26 | 0.791 |
| retention - free choice 1 | rl | 0.07 | 0.07 | Inf | -0.07-0.21 | 0.96 | 0.339 |
| retention - free choice 2 | rl | 0.05 | 0.07 | Inf | -0.09-0.19 | 0.72 | 0.469 |
| transfer - free choice 1 | rl | 0.18 | 0.07 | Inf | 0.04-0.32 | 2.58 | 0.010 |
| transfer - free choice 2 | rl | 0.16 | 0.07 | Inf | 0.03-0.30 | 2.34 | 0.019 |
| transfer - retention | rl | 0.11 | 0.08 | Inf | -0.04-0.27 | 1.42 | 0.155 |
| free choice 2 - free choice 1 | slfc | 0.08 | 0.06 | Inf | -0.04-0.21 | 1.32 | 0.188 |
| retention - free choice 1 | slfc | 0.31 | 0.07 | Inf | 0.18-0.44 | 4.59 | < 0.001 |
| retention - free choice 2 | slfc | 0.23 | 0.07 | Inf | 0.09-0.36 | 3.31 | < 0.001 |
| transfer - free choice 1 | slfc | 0.36 | 0.07 | Inf | 0.23-0.50 | 5.33 | < 0.001 |
| transfer - free choice 2 | slfc | 0.28 | 0.07 | Inf | 0.14-0.42 | 4.02 | < 0.001 |
| transfer - retention | slfc | 0.05 | 0.06 | Inf | -0.07-0.18 | 0.85 | 0.396 |

Formula: glmer(chosetheorybest ∼ treatment * session + (1 | skigroup/skier).

rl = reinforcement learning; slfc = supervised (free choice) learning

**Supplementary Table 4**: Predicted probability difference between groups

| Contrast | Session | Estimate | SE | df | CI | z | p |
|---|---|---|---|---|---|---|---|
| rl - slfc | free choice 1 | 0.01 | 0.13 | Inf | -0.24-0.27 | 0.11 | 0.911 |
| rl - slfc | free choice 2 | -0.05 | 0.13 | Inf | -0.31-0.20 | -0.39 | 0.693 |
| rl - slfc | retention | -0.23 | 0.13 | Inf | -0.48-0.03 | -1.75 | 0.079 |
| rl - slfc | transfer | -0.17 | 0.12 | Inf | -0.40-0.07 | -1.41 | 0.159 |

Formula: glmer(chosetheorybest ∼ treatment * session + (1 | skigroup/skier).

rl = reinforcement learning; slfc = supervised (free choice) learning.

**Supplementary Table 5**: Strategy choice: Predicted probability change in probability of choosing the individual skier's estimated best strategy

| Contrast | Treatment | Estimate | SE | df | CI | z | p |
|---|---|---|---|---|---|---|---|
| free choice 2 - free choice 1 | rl | 0.20 | 0.06 | Inf | 0.09-0.32 | 3.39 | $< 0.001$ |
| retention - free choice 1 | rl | 0.13 | 0.06 | Inf | 0.00-0.26 | 2.02 | 0.043 |
| retention - free choice 2 | rl | -0.07 | 0.06 | Inf | -0.19-0.04 | -1.25 | 0.213 |
| transfer - free choice 1 | rl | 0.21 | 0.06 | Inf | 0.09-0.34 | 3.30 | $< 0.001$ |
| transfer - free choice 2 | rl | 0.01 | 0.05 | Inf | -0.09-0.11 | 0.18 | 0.854 |
| transfer - retention | rl | 0.08 | 0.06 | Inf | -0.04-0.20 | 1.31 | 0.190 |
| free choice 2 - free choice 1 | slfc | 0.09 | 0.06 | Inf | -0.03-0.21 | 1.48 | 0.140 |
| retention - free choice 1 | slfc | 0.21 | 0.07 | Inf | 0.08-0.34 | 3.11 | 0.002 |
| retention - free choice 2 | slfc | 0.12 | 0.06 | Inf | -0.00-0.24 | 1.88 | 0.060 |
| transfer - free choice 1 | slfc | 0.23 | 0.07 | Inf | 0.10-0.36 | 3.39 | $< 0.001$ |
| transfer - free choice 2 | slfc | 0.14 | 0.06 | Inf | 0.02-0.26 | 2.21 | 0.027 |
| transfer - retention | slfc | 0.02 | 0.06 | Inf | -0.09-0.13 | 0.35 | 0.727 |

Formula: glmer(choseestimatedbest treatment * session + (1 | skigroup/skier).

rl = reinforcement learning; slfc = supervised (free choice) learning

**Supplementary Table 6**: Strategy choice: Predicted probability difference of choosing the individual skier's estimated best strategy at each session

| Contrast | Session | Estimate | SE | df | CI | z | p |
|---|---|---|---|---|---|---|---|
| rl - slfc | block2 | 0.01 | 0.12 | Inf | -0.22-0.24 | 0.12 | 0.904 |
| rl - slfc | block3 | 0.13 | 0.10 | Inf | -0.06-0.32 | 1.30 | 0.194 |
| rl - slfc | retention | -0.06 | 0.10 | Inf | -0.26-0.14 | -0.61 | 0.544 |
| rl - slfc | transfer | 0.00 | 0.09 | Inf | -0.17-0.17 | 0.00 | 0.999 |

Formula: glmer(choseestimatedbest $\sim$ treatment * session + (1 | skigroup/skier).

rl = reinforcement learning; slfc = supervised (free choice) learning

**Supplementary Table 7**: Strategy evaluation: Estimated differences between strategy ranking during familiarization

| Term | Estimate | SE | CI | t | p |
|---|---|---|---|---|---|
| (Intercept) | 2.50 | 0.03 | 2.44-2.56 | 85.64 | < 0.001 |
| slfc | 0.00 | 0.08 | -0.16-0.16 | 0.00 | 1.000 |
| slts | 0.00 | 0.05 | -0.11-0.11 | 0.00 | 1.000 |
| ranktime | 0.00 | 0.01 | -0.02-0.02 | 0.00 | 1.000 |
| rl : b | -1.72 | 0.11 | -1.94–1.50 | -15.47 | < 0.001 |
| slfc : b | -1.43 | 0.19 | -1.81–1.05 | -7.38 | < 0.001 |
| slts : b | -1.20 | 0.11 | -1.41–0.99 | -11.32 | < 0.001 |
| rl : c | -1.05 | 0.11 | -1.27–0.83 | -9.42 | < 0.001 |
| slfc : c | -0.70 | 0.19 | -1.08–0.32 | -3.60 | < 0.001 |
| slts : c | -0.95 | 0.11 | -1.16–0.74 | -8.94 | < 0.001 |
| rl : d | -2.06 | 0.11 | -2.28–1.84 | -18.49 | < 0.001 |
| slfc : d | -2.09 | 0.19 | -2.47–1.71 | -10.77 | < 0.001 |
| slts : d | -2.41 | 0.11 | -2.61–2.20 | -22.67 | < 0.001 |
| slfc : ranktime | -0.00 | 0.02 | -0.04-0.04 | -0.00 | 1.000 |
| slts : ranktime | -0.00 | 0.02 | -0.04-0.04 | -0.00 | 1.000 |
| rl : b : ranktime | -0.15 | 0.04 | -0.22–0.07 | -3.95 | < 0.001 |
| slfc : b : ranktime | -0.12 | 0.05 | -0.22–0.02 | -2.45 | 0.014 |
| slts : b : ranktime | -0.13 | 0.04 | -0.20–0.06 | -3.65 | < 0.001 |
| rl : c : ranktime | 0.05 | 0.04 | -0.03-0.12 | 1.21 | 0.225 |
| slfc : c : ranktime | -0.01 | 0.05 | -0.11-0.09 | -0.27 | 0.791 |
| slts : c : ranktime | -0.06 | 0.04 | -0.13-0.01 | -1.78 | 0.076 |
| rl : d : ranktime | -0.09 | 0.04 | -0.17–0.02 | -2.51 | 0.012 |
| slfc : d : ranktime | -0.09 | 0.05 | -0.19-0.01 | -1.85 | 0.065 |
| slts : d : ranktime | -0.15 | 0.04 | -0.22–0.08 | -4.18 | < 0.001 |

Formula: lm(value ∼ treatment/strategy * ranktime)

rl = reinforcement learning; slfc = supervised (free choice) learning, slts = supervised (target skill) learning, b="extend", c="rock skis forward", d="extend with rock skis forward"

**Supplementary Table 8**: Strategy evaluation: Estimated differences between groups on strategy ranking during familiarization

| Term | Estimate | SE | CI | t | p |
|---|---|---|---|---|---|
| (Intercept) | 2.50 | 0.03 | 2.44-2.56 | 85.64 | $< 0.001$ |
| b | -1.45 | 0.08 | -1.61–1.29 | -17.58 | $< 0.001$ |
| c | -0.90 | 0.08 | -1.06–0.74 | -10.88 | $< 0.001$ |
| d | -2.18 | 0.08 | -2.35–2.02 | -26.46 | $< 0.001$ |
| ranktime | -0.00 | 0.01 | -0.02-0.02 | -0.00 | 1.000 |
| a : slfc | -0.15 | 0.16 | -0.46-0.16 | -0.96 | 0.340 |
| b : slfc | 0.14 | 0.16 | -0.17-0.45 | 0.87 | 0.385 |
| c : slfc | 0.20 | 0.16 | -0.11-0.51 | 1.26 | 0.209 |
| d : slfc | -0.19 | 0.16 | -0.50-0.13 | -1.17 | 0.242 |
| a : slts | -0.07 | 0.11 | -0.28-0.15 | -0.63 | 0.531 |
| b : slts | 0.45 | 0.11 | 0.24-0.67 | 4.17 | $< 0.001$ |
| c : slts | 0.03 | 0.11 | -0.18-0.24 | 0.29 | 0.772 |
| d : slts | -0.42 | 0.11 | -0.63–0.20 | -3.83 | $< 0.001$ |
| b : ranktime | -0.13 | 0.02 | -0.18–0.09 | -5.56 | $< 0.001$ |
| c : ranktime | -0.01 | 0.02 | -0.06-0.04 | -0.43 | 0.665 |
| d : ranktime | -0.11 | 0.02 | -0.16–0.06 | -4.65 | $< 0.001$ |
| a : slfc : ranktime | 0.01 | 0.04 | -0.08-0.10 | 0.19 | 0.847 |
| b : slfc : ranktime | 0.03 | 0.04 | -0.05-0.12 | 0.73 | 0.465 |
| c : slfc : ranktime | -0.05 | 0.04 | -0.14-0.04 | -1.13 | 0.259 |
| d : slfc : ranktime | 0.01 | 0.04 | -0.08-0.10 | 0.21 | 0.837 |
| a : slts : ranktime | 0.04 | 0.04 | -0.03-0.11 | 1.00 | 0.317 |
| b : slts : ranktime | 0.05 | 0.04 | -0.02-0.13 | 1.48 | 0.140 |
| c : slts : ranktime | -0.07 | 0.04 | -0.14–0.00 | -1.97 | 0.049 |
| d : slts : ranktime | -0.02 | 0.04 | -0.09-0.05 | -0.50 | 0.615 |

Formula: lm(value ∼ strategy / treatment * ranktime)

rl = reinforcement learning; slfc = supervised (free choice) learning, slts = supervised (target skill) learning, a="stand against", "b="extend", c="rock skis forward", d="extend with rock skis forward"

**Supplementary Table 9**: Strategy evaluation: Estimated slope for each strategy across session

| Term | Estimate | SE | CI | t | p |
|---|---|---|---|---|---|
| (Intercept) | 2.50 | 0.03 | 2.44-2.56 | 85.64 | $< 0.001$ |
| slfc | 0.00 | 0.08 | -0.16-0.16 | 0.00 | 1.000 |
| slts | 0.00 | 0.05 | -0.11-0.11 | 0.00 | 1.000 |
| b | -1.45 | 0.08 | -1.61–1.29 | -17.58 | $< 0.001$ |
| c | -0.90 | 0.08 | -1.06–0.74 | -10.88 | $< 0.001$ |
| d | -2.18 | 0.08 | -2.35–2.02 | -26.46 | $< 0.001$ |
| slfc : b | 0.29 | 0.22 | -0.15-0.73 | 1.29 | 0.197 |
| slts : b | 0.52 | 0.15 | 0.22-0.82 | 3.39 | $< 0.001$ |
| slfc : c | 0.35 | 0.22 | -0.09-0.79 | 1.56 | 0.118 |
| slts : c | 0.10 | 0.15 | -0.20-0.40 | 0.65 | 0.517 |
| slfc : d | -0.03 | 0.22 | -0.47-0.41 | -0.15 | 0.879 |
| slts : d | -0.35 | 0.15 | -0.65–0.05 | -2.26 | 0.024 |
| rl : a : ranktime | 0.05 | 0.03 | -0.00-0.10 | 1.86 | 0.064 |
| slfc : a : ranktime | 0.06 | 0.04 | -0.01-0.13 | 1.61 | 0.107 |
| slts : a : ranktime | 0.09 | 0.03 | 0.04-0.13 | 3.40 | $< 0.001$ |
| rl : b : ranktime | -0.10 | 0.03 | -0.15–0.05 | -3.73 | $< 0.001$ |
| slfc : b : ranktime | -0.07 | 0.04 | -0.14-0.00 | -1.85 | 0.065 |
| slts : b : ranktime | -0.04 | 0.03 | -0.09-0.00 | -1.77 | 0.077 |
| rl : c : ranktime | 0.09 | 0.03 | 0.04-0.15 | 3.57 | $< 0.001$ |
| slfc : c : ranktime | 0.04 | 0.04 | -0.03-0.11 | 1.24 | 0.216 |
| slts : c : ranktime | 0.02 | 0.03 | -0.03-0.07 | 0.89 | 0.376 |
| rl : d : ranktime | -0.04 | 0.03 | -0.10-0.01 | -1.70 | 0.090 |
| slfc : d : ranktime | -0.04 | 0.04 | -0.11-0.03 | -1.00 | 0.317 |
| slts : d : ranktime | -0.06 | 0.03 | -0.11–0.01 | -2.51 | 0.012 |

Formula: lm(value ∼ treatment * strategy / ranktime)

rl = reinforcement learning; slfc = supervised (free choice) learning, slts = supervised (target skill) learning, a="stand against", b="extend", c="rock skis forward", d="extend with rock skis forward"

**Supplementary Table 10**: Strategy outcome: Estimated difference between strategies during forced exploration

| Term | Estimate | SE | df | CI | t | p |
|---|---|---|---|---|---|---|
| (Intercept) | 16.88 | 0.11 | 3.99 | 16.56-17.20 | 147.94 | $< 0.001$ |
| supervised (free choice) | 0.06 | 0.13 | 91.87 | -0.21-0.32 | 0.42 | 0.674 |
| supervised (target skill) | 0.12 | 0.13 | 91.72 | -0.14-0.37 | 0.90 | 0.370 |
| reinforcement learning : c-a | -0.22 | 0.03 | 641.78 | -0.29–0.15 | -6.39 | $< 0.001$ |
| supervised (free choice) : c-a | -0.16 | 0.03 | 641.98 | -0.23–0.10 | -4.80 | $< 0.001$ |
| supervised (target skill) : c-a | -0.22 | 0.03 | 641.72 | -0.28–0.15 | -6.53 | $< 0.001$ |
| reinforcement learning : b-c | -0.22 | 0.03 | 641.75 | -0.29–0.16 | -6.47 | $< 0.001$ |
| supervised (free choice) : b-c | -0.15 | 0.03 | 641.93 | -0.22–0.09 | -4.51 | $< 0.001$ |
| supervised (target skill) : b-c | -0.23 | 0.03 | 641.75 | -0.30–0.17 | -7.00 | $< 0.001$ |
| reinforcement learning : d-b | 0.00 | 0.03 | 641.72 | -0.07-0.07 | 0.02 | 0.984 |
| supervised (free choice) : d-b | -0.05 | 0.03 | 641.97 | -0.12-0.01 | -1.53 | 0.125 |
| supervised (target skill) : d-b | -0.01 | 0.03 | 641.80 | -0.07-0.06 | -0.21 | 0.834 |
| sd(Intercept) | 0.52 | | | | | |
| sd(Intercept) | 0.20 | | | | | |
| sd(Observation) | 0.19 | | | | | |

Formula: lmer(racingtime ∼ treatment / strategy + (1 — skigroup / skier)

a="stand against", b="extend", c="rock skis forward", d="extend with rock skis forward"

For this model we used a sliding difference contrast coding scheme

**Supplementary Table 11**: Strategy outcome: Estimated difference between groups per strategy

| Term | Estimate | SE | df | CI | t | p |
|---|---|---|---|---|---|---|
| (Intercept) | 16.88 | 0.11 | 3.99 | 16.56-17.20 | 147.94 | $< 0.001$ |
| b | -0.40 | 0.02 | 641.79 | -0.44–0.37 | -20.63 | $< 0.001$ |
| c | -0.20 | 0.02 | 641.83 | -0.24–0.16 | -10.22 | $< 0.001$ |
| d | -0.42 | 0.02 | 641.78 | -0.46–0.38 | -21.72 | $< 0.001$ |
| a : supervised (free choice) | -0.01 | 0.13 | 101.33 | -0.28-0.26 | -0.07 | 0.945 |
| b : supervised (free choice) | 0.12 | 0.13 | 101.55 | -0.15-0.39 | 0.87 | 0.386 |
| c : supervised (free choice) | 0.05 | 0.14 | 101.61 | -0.22-0.32 | 0.36 | 0.716 |
| d : supervised (free choice) | 0.06 | 0.13 | 101.04 | -0.20-0.33 | 0.48 | 0.632 |
| a : supervised (target skill) | 0.17 | 0.13 | 101.01 | -0.09-0.44 | 1.28 | 0.203 |
| b : supervised (target skill) | 0.17 | 0.13 | 100.98 | -0.10-0.44 | 1.26 | 0.209 |
| c : supervised (target skill) | 0.18 | 0.13 | 101.24 | -0.09-0.45 | 1.33 | 0.188 |
| d : supervised (target skill) | 0.16 | 0.13 | 101.33 | -0.11-0.43 | 1.21 | 0.231 |
| sd(Intercept) | 0.52 | | | | | |
| sd(Intercept) | 0.20 | | | | | |
| sd(Observation) | 0.19 | | | | | |

Formula: lmer(racingtime ~ strategy / treatment + (1 | skigroup / skier)

a="stand against", b="extend", c="rock skis forward", d="extend with rock skis forward"

**Supplementary Table 12**: Strategy outcome: Estimated change on "extend with rock skis forward"

| Term | Estimate | SE | df | CI | t | p |
|---|---|---|---|---|---|---|
| (Intercept) | 16.59 | 0.16 | 6.81 | 16.20-16.98 | 101.76 | $< 0.001$ |
| supervised (free choice) | 0.08 | 0.13 | 95.97 | -0.18-0.35 | 0.64 | 0.522 |
| supervised (target skill) | 0.06 | 0.13 | 94.98 | -0.20-0.32 | 0.44 | 0.660 |
| reinforcement learning : session number | -0.09 | 0.01 | 1065.01 | -0.11–0.07 | -8.15 | $< 0.001$ |
| supervised (free choice) : session number | -0.10 | 0.01 | 1063.28 | -0.11–0.08 | -10.01 | $< 0.001$ |
| supervised (target skill) : session number | -0.08 | 0.01 | 1058.03 | -0.09–0.06 | -9.25 | $< 0.001$ |
| sd(Intercept) | 0.51 | | | | | |
| sd(Intercept) | 0.27 | | | | | |
| sd(Observation) | 0.18 | | | | | |

Formula: lmer(racetime ~ treatment / sessionnumber + (1 | skigroup / skier))

rl = reinforcement learning; slfc = supervised (free choice) learning, slts = supervised (target skill) learning, a="stand against", b="extend", c="rock skis forward", d="extend with rock skis forward"

**Supplementary Table 13**: Strategy outcome: Estimated change for each strategy

| Term | Estimate | SE | df | CI | t | p |
|---|---|---|---|---|---|---|
| (Intercept) | 16.80 | 0.11 | 4.05 | 16.48-17.12 | 146.23 | < 0.001 |
| slfc | 0.06 | 0.13 | 60.23 | -0.21-0.32 | 0.44 | 0.661 |
| b | -0.44 | 0.02 | 1378.01 | -0.48–0.40 | -20.15 | < 0.001 |
| c | -0.21 | 0.02 | 1378.03 | -0.25–0.16 | -8.82 | < 0.001 |
| d | -0.45 | 0.02 | 1378.05 | -0.50–0.41 | -21.04 | < 0.001 |
| slfc : b | 0.17 | 0.04 | 1378.01 | 0.09-0.26 | 3.98 | < 0.001 |
| slfc : c | 0.07 | 0.05 | 1378.03 | -0.02-0.16 | 1.47 | 0.142 |
| slfc : d | 0.08 | 0.04 | 1378.05 | -0.00-0.17 | 1.95 | 0.051 |
| rl : a : session number | 0.01 | 0.05 | 1378.53 | -0.09-0.10 | 0.18 | 0.860 |
| slfc : a : session number | -0.07 | 0.03 | 1378.40 | -0.13–0.01 | -2.36 | 0.018 |
| rl : b : session number | -0.11 | 0.01 | 1378.80 | -0.13–0.09 | -9.80 | < 0.001 |
| slfc : b : session number | -0.07 | 0.01 | 1379.01 | -0.10–0.05 | -5.89 | < 0.001 |
| rl : c : session number | -0.12 | 0.03 | 1380.31 | -0.17–0.07 | -4.58 | < 0.001 |
| slfc : c : session number | -0.13 | 0.02 | 1378.76 | -0.17–0.09 | -6.13 | < 0.001 |
| rl : d : session number | -0.11 | 0.01 | 1378.82 | -0.13–0.08 | -9.49 | < 0.001 |
| slfc : d : session number | -0.10 | 0.01 | 1378.28 | -0.12–0.08 | -10.59 | < 0.001 |
| sd(Intercept) | 0.52 | | | | | |
| sd(Intercept) | 0.19 | | | | | |
| sd(Observation) | 0.19 | | | | | |

Formula: lmer(racetime ∼ treatment * strategy / sessionnumber + (1 | skigroup / skier))

rl = reinforcement learning; slfc = supervised (free choice) learning, a="stand against", b="extend", c="rock skis forward", d="extend with rock skis forward"

**Supplementary Table 14**: Strategy outcome: Estimated difference in change between groups for each strategy

| Term | Estimate | SE | df | CI | t | p |
|---|---|---|---|---|---|---|
| (Intercept) | 16.80 | 0.11 | 4.05 | 16.48-17.12 | 146.23 | < 0.001 |
| b | -0.44 | 0.02 | 1378.01 | -0.48–0.40 | -20.15 | < 0.001 |
| c | -0.21 | 0.02 | 1378.03 | -0.25–0.16 | -8.82 | < 0.001 |
| d | -0.45 | 0.02 | 1378.05 | -0.50–0.41 | -21.04 | < 0.001 |
| session number | -0.09 | 0.01 | 1378.67 | -0.10–0.07 | -9.99 | < 0.001 |
| a : slfc | -0.02 | 0.14 | 66.50 | -0.30-0.25 | -0.17 | 0.866 |
| b : slfc | 0.15 | 0.14 | 64.08 | -0.12-0.42 | 1.12 | 0.268 |
| c : slfc | 0.05 | 0.14 | 65.91 | -0.23-0.32 | 0.33 | 0.740 |
| d : slfc | 0.06 | 0.13 | 63.63 | -0.21-0.33 | 0.45 | 0.653 |
| b : session number | -0.06 | 0.03 | 1378.48 | -0.12–0.00 | -1.98 | 0.047 |
| c : session number | -0.09 | 0.03 | 1378.72 | -0.16–0.03 | -2.78 | 0.005 |
| d : session number | -0.07 | 0.03 | 1378.41 | -0.13–0.01 | -2.44 | 0.015 |
| a : slfc : session number | -0.08 | 0.06 | 1378.54 | -0.19-0.03 | -1.40 | 0.163 |
| b : slfc : session number | 0.04 | 0.02 | 1378.88 | 0.01-0.07 | 2.34 | 0.020 |
| c : slfc : session number | -0.01 | 0.03 | 1379.85 | -0.07-0.06 | -0.23 | 0.819 |
| d : slfc : session number | 0.00 | 0.01 | 1378.56 | -0.03-0.03 | 0.20 | 0.842 |
| sd(Intercept) | 0.52 | | | | | |
| sd(Intercept) | 0.19 | | | | | |
| sd(Observation) | 0.19 | | | | | |

Formula: lmer(racetime ∼ strategy / treatment * sessionnumber + (1 | skigroup / skier))

rl = reinforcement learning; slfc = supervised (free choice) learning, a="stand against", b="extend", c="rock skis forward", d="extend with rock skis forward"

### Supplementary Methods A: Snow preparation

We dedicated a substantial amount of time and effort to prepare the hill for our learning experiment. Our primary objective was to ensure that the snow conditions were as identical and fair as possible for all participants. To achieve this, we collaborated closely with the SNØ facility's staff and the coaches of the Norwegian Alpine Ski team. Together, we devised a comprehensive plan to achieve consistency before each group of skiers took to the slopes. This report will detail the steps we took to ensure the snow was ready for our skiers and provide insight into the reasoning behind our choices. The purpose of this supplementary note is to assist you in evaluating our study and to transparently document our commitment to delivering the best possible conditions for our skiers.

#### Ski group A

About two weeks before testing Group A, new snow was created on the racing hill. The evening before data collection, machines groomed the racing hill (Fig. 1a:left). We then watered the hill and left it to freeze overnight to create a hard and firm surface (Fig. 1a:right).

#### Ski group B

Ski group B began testing the day after ski group A completed their testing. Once group A finished, we inspected the race hill(Fig. 1b:left), noticed some small holes, and decided to fill them before watering it again. (Fig. 1b:right).

#### Ski group C

After ski group B, the race hill started getting icy, lacking grip in some areas. We were concerned that watering it again might worsen conditions, making it too challenging for skiers. Consequently, we opted to gently groom the hill with a machine and let the grooves set for a couple of days. See Fig. 1c for the result of this process.

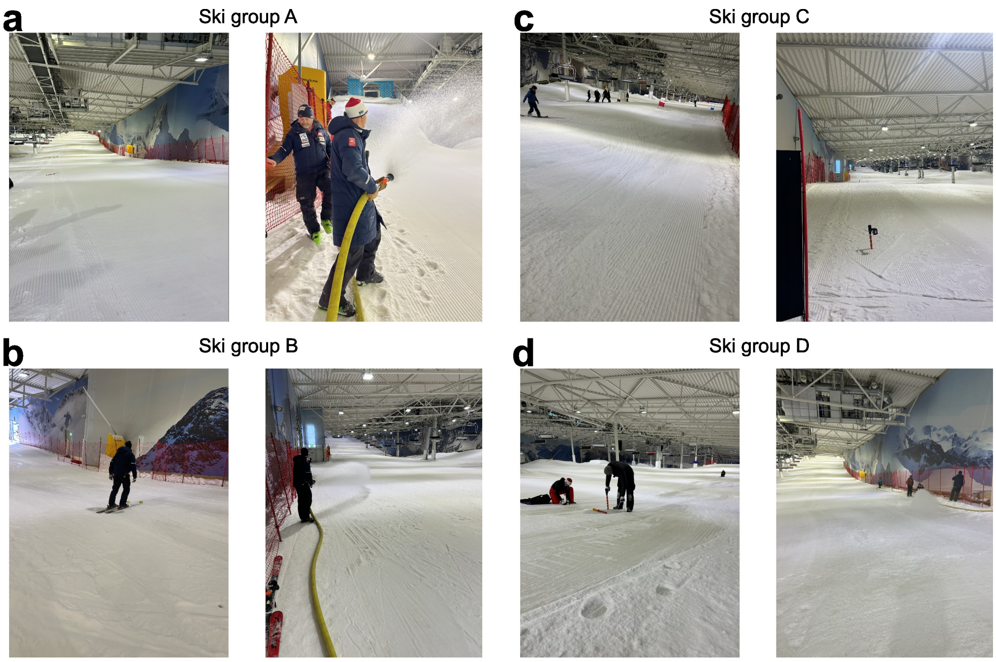


**Fig. 1**: Images showing the hill preparation for the four ski groups. **a**. Shows the racing hill for ski group A. The left image displays the racing hill the evening before data collection after it was groomed. The right image shows the watering process for ski group A. **b**. Shows the racing hill for ski group B. The left image depicts the course inspection immediately after ski group A finished their testing. The right image shows the watering process for ski group B. Note that we did not groom the course this time, so in the image, you can see some uneven surfaces on the snow, which were evened out by watering it. **c**. Shows the racing hill for ski group C. The left image displays the hill after it was groomed and left overnight. The right image shows the same but from the bottom. **d**. Shows the racing hill for ski group D. The left image illustrates the hill for ski group C on their retention test. Note the icy surface, which was the reason why we produced new snow. The right image shows the watering process for ski group D.

#### Ski group D

After Ski Group C, the race hill needed new snow because some areas had become icy, with minimal grip (Fig. reffig:snowprepd:left)). Although the conditions were suitable for Ski Group C, they would not have worked for a new ski group undergoing testing. Consequently, we decided to produce new snow two days before Ski Group D started their training. This fresh snow was pushed into the racing hill the day before testing and groomed. Subsequently, we watered the hill and let it freeze overnight (Fig. 1d:right).

### Supplementary Methods B: Course setting

We used a standard procedure to set the slalom courses, ensuring a fixed length and offset. First, we stretched a taut rope between two nails on either side of the ski hill. This rope helped us locate the exact starting line consistently from day to day. From the nail on the skier's right, we measured 6 meters into the slope. We did the same from a fixed point approximately 50 meters down the course, but here we measured 3.4 meters out. Then, we pulled a 50-meter-long measuring tape between the two points to establish the line down the hill. We chose a measuring tape over a rope because ropes tend to expand and contract when they get wet and dry, respectively (see Fig.2a for an image illustrating this process).

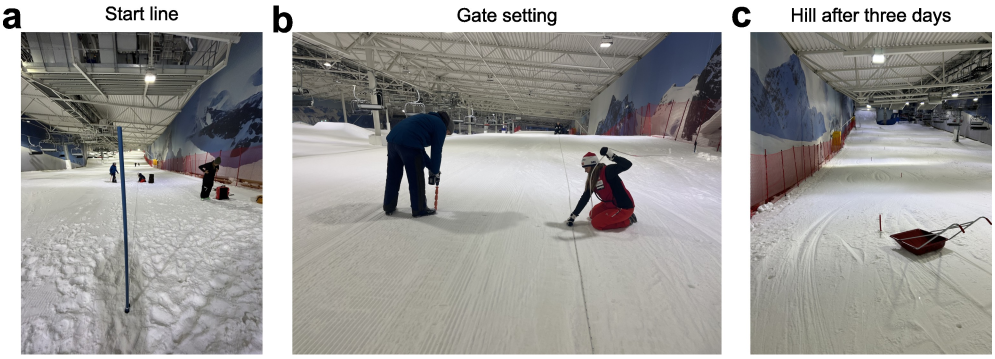


**Fig. 2**: Images showing the procedure for course setting. **a**. The image illustrates the process of establishing the straight reference line down the hill. The long blue gate is positioned 6 meters from the nail on the skier's right. **b**. This image showcases the gate-setting method. A white rope is secured to the reference line with a carabiner hook, and a marker on the rope indicates a distance of 1.9 meters for placing the gate. **c**. The image reveals the tracks left behind by a group of skiers that was tested

Once the 50-meter straight line was established, we laid out a rope segment attached to a carabiner hooked onto the long rope. We moved this segment 1.9 meters out from the rope in one turn and 0 meters in the other, with a 10-meter vertical space. This ensured that the course followed a straight line down the slope and was set with the correct offset (see Fig. 2b for an image illustrating this process). Once all the gates along the 50-meter measurement tape were set, we used a new fixation point down the slope and continued down the course. We practiced this procedure several times before the experiment, and the variation in course setting was a maximum of 10 cm at the end of the course.

Due to wear and tear on the trails, we opted to shift the course laterally or rotate it, depending on the situation. With the exception of one skiing group, we followed the following practice. On day 1, we set the course as described above. On day 2, we rotated the course so that the first turn went in the opposite direction of day 1. On the third day, we shifted the course closer to the wall on skiers' right (see Fig. 2c for an image showing the tracks in the hill left after a skigroup had completed the experiment).

### Supplementary Methods C: Description of strategies

This supplementary describes each strategy in more detail. The "stand against" strategy emphasized maintaining a stable stance against external forces without body extension along the body's longitudinal axis or rocking skis forward. This term is frequently used by Norwegian ski coaches when communicating with skiers to help them improve their race times.

The "rock skis forward" strategy involves rocking the ski forward during the turning phase. This action effectively regulates the distribution of pressure over the skis. During the initiation and control phases of a turn, the pressure is generally shifted forward to bend the ski's forebody, increasing friction with the snow and enabling it to turn more sharply. However, at some point, during turn progression, the skier aims to stop turning and therefore shifts the pressure further back on the skis, thereby reducing turning and braking forces [1, 2]. Investigations of elite alpine ski racers have shown that high performing skiers tend to rock skis more forward and pressure the back part of the ski for considerable longer time during a turn, than slower skiers [3]. To make the information more specific for the skiers, we communicated, that the maximum range of the rocking movement was about 30-50 centimeters from gate passage to completion of the turn, which is in correspondence with biomechanical evidence of elite ski racers [3].

The "extend" strategy involves extending the body from a laterally tilted position during the turn, closer to the turn's center of rotation. When skiers move their center of mass this way by extending their legs, they increase their kinetic rotational energy under certain situations. According to Lind and Sanders [4] model, the skier achieves this effect by shortening the radius of the axis about which they rotate, which will reduce the moment of inertia and consequently increase the rotational kinetic energy of the system under the assumption that angular momentum is conserved. In their model, the gain in rotational kinetic energy from this motion is proportional to the amount of work the skier does against the centrifugal force (from the skier's frame of reference); therefore, a larger extension movement will accomplish a greater increase in rotational kinetic energy. Simulation studies of individuals extending their body in rollers or during carved slalom turns have shown that this movement can increase speed [5, 6], and the effect has been observed in various ways with elite skiers during training and competition [3, 7, 8].

Finally, "extend with rocking skis forward" was expected to be the best strategy combining the two effects from extending and rocking skis forward, and we therefore defined this as the theoretical best strategy. Simulations of skiers extending their bodies in the bottom of a roller have observed an additional effect of rocking the skis forward [5]

### Supplementary Methods D: Coach setup

We set up three coach stations in the finishing area for the supervised learning groups, one for each coach. The space between the coaches was approximately 3 to 5 meters. We used wall dividers to prevent information leakage between the coaches. In addition, the background noise in the ski hall was generally high, and it was difficult to perceive information without standing close to the person. The supervised (target skill) learning coach was behind the two other coaches to prevent the other coaches from seeing what he did. Each coach had a monitor where they could see their own (but not the other) skiers' race.

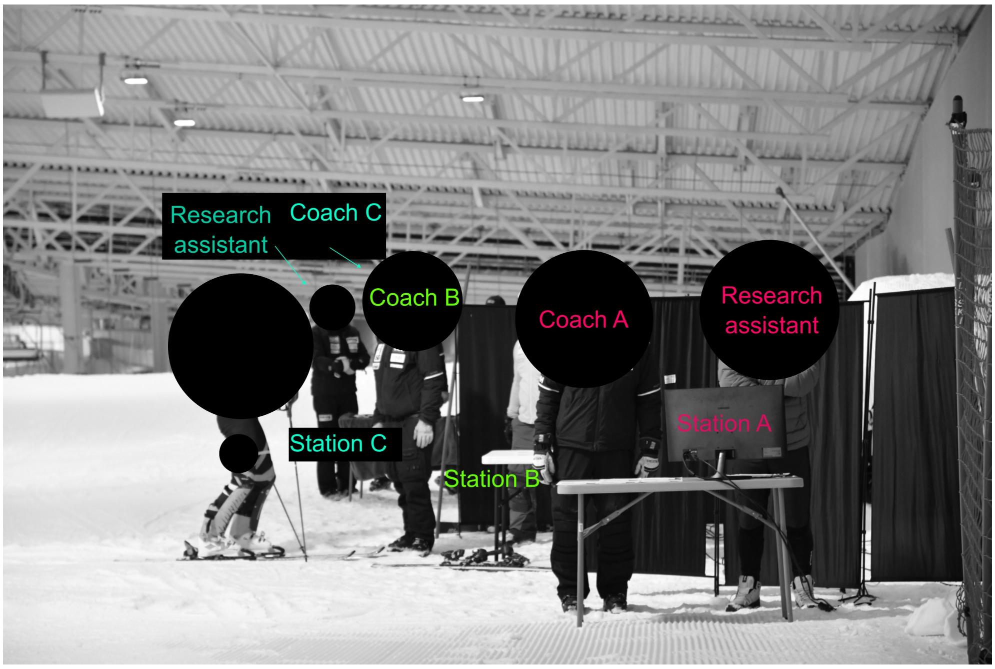


**Fig. 3**: Images shows the setup in the finish area

#### Supplementary Discussion

To convert race time to FIS World Ranking, we assumed that each second corresponded to a 7-point FIS. We could then multiply by 0.12 * 7 to find the difference in FIS points. Then, we used the median rank in our sample and calculated what this difference corresponded to in the World Ranking. We performed this analysis for both females and men. Due to confidentiality, we do not want to say which FIS list we used for this conversion. If the reader thinks 7 points is too small or large, then we welcome the reader to change this number up or down. We only used this conversion to help readers evaluating the effect.
